# Temperature influences West Nile virus evolution and adaptation

**DOI:** 10.1101/2025.02.28.640855

**Authors:** R.L. Fay, M. Cruz-Loya, J.G. Maffei, E.A. Mordecai, A.T Ciota

**Affiliations:** Department of Biomedical Sciences, State University of New York at Albany School of Public Health, Rensselaer, NY, USA; The Arbovirus Laboratory, Wadsworth Center, New York State Department of Health, Slingerlands, NY, USA; Biology Department, Stanford University, Stanford, CA, USA

**Keywords:** West Nile virus, Temperature, Vector-borne, Evolution, Adaptation

## Abstract

West Nile virus (WNV), the most common mosquito-borne disease in the continental U.S., is vectored by *Culex* spp. mosquitoes. Since its introduction to New York State (NYS) in 1999, WNV has become endemic. NYS temperatures have risen by 0.14°C per decade since 1900, with larger increases linked to higher WNV transmission. Using surveillance and sequencing data, we find a significant correlation between rising temperatures, increased WNV genetic diversity, and higher prevalence. Given the experimentally demonstrated role of temperature influencing WNV fitness, we hypothesized that contemporary strains should exhibit greater fitness in mosquitoes at higher temperatures compared to historic strains. To test this, we analyzed genetically distinct WNV strains from mosquitoes collected during recent warm summers (2017 and 2018) and cooler historic summers (2003 and 2004). Assessing *Culex pipiens* vector competence and calculating the relative R₀ at 20°C, 24°C, and 28°C, we found that contemporary strains exhibit higher transmission potential at increased temperatures. Our results show that contemporary WNV strains possess greater phenotypic and genotypic diversity, facilitating the emergence of strains with enhanced transmission potential in a warming climate.

## Introduction

West Nile virus (WNV) is a vector-borne pathogen, primarily transmitted by *Culex* species mosquitoes. WNV emerged in the Western Hemisphere in 1999 in New York State (NYS) and was linked to an outbreak of neuroinvasive illnesses[1–3]. Since then, WNV has spread across the United States (US)[4]. WNV is a member of the *Flavivirdae* family, a group of single-stranded positive-sense RNA genome viruses including Zika virus (ZIKV), dengue virus (DENV), yellow fever virus (YFV), and several other human pathogens[5]. A mosquito acquires WNV when it takes a blood meal from an infected bird[6]. The virus then replicates in the midgut of the mosquito, escapes the gut to the hemolymph, infects and replicates in the salivary glands, and is transmitted through saliva during the mosquito’s next blood meal[2,7]. Previous studies have demonstrated a relationship between increases in environmental temperature and increased WNV vector competence as well as accelerated extrinsic incubation period (EIP)[8–10].

Across the globe, average temperatures are increasing[11], with consequences for vector-borne disease transmission. In NYS, temperatures have risen at an average rate of 0.14°C per decade since 1900 and are predicted to increase more than 2.28°C by 2080[12]. Mosquitoes and the pathogens they transmit are ectothermic organisms that rely on external sources of heat for temperature regulation. Such organisms are susceptible to temperature fluctuations, which can alter mosquito life history traits, viral replication, and pathogen transmissibility. Most physiological and life history traits, including those of WNV and its vectors, respond unimodally to temperature, with a thermal minimum, optimum, and maximum[13].

WNV can adapt rapidly to changing environments. The virus circulates between its mosquito and avian hosts, experiencing different thermal environments throughout replication[14]. In the last 25 years, it has become apparent that WNV evolution plays a key role in disease transmission. Previous studies demonstrated that the WN02 genotype, which displaced the ancestral NY99 genotype, acquired adaptive mutations that increased transmission by *Culex* mosquitoes and contributed to the spread of WNV across America[15]. Further, the adaptive advantage of WN02 strains is enhanced at higher temperatures[8]. In more recent years, new strains of WNV associated with increased prevalence and further gains in transmissibility have emerged, yet the thermal sensitivity of contemporary strains has yet to be evaluated[16]. Temperature has been shown to drive WNV diversification over time in North America[17]. In addition, previous work has shown that warmer temperature increases viral replication and mutation frequency *in vitro*[18]. Specifically, WNV passaged in mosquito cells at 30°C were more broadly adapted and acquired more nonsynonymous mutations when compared to virus passaged at 25°C[18].

We hypothesized that temperature increases in the Northeast U.S. since the emergence of WNV have influenced virus diversification, adaptation, and alterations in thermal sensitivity[17,19]. We asked three main questions: 1) Does intraseason diversity of WNV increase with time and temperature? 2) What is the extent of strain-specific variability in temperature-dependent transmission of WNV? and 3) Does evolution under higher temperatures drive temperature-dependent adaptation? To address these questions, we retrospectively determined the relationship between annual WNV amino acid diversity and mean temperature within the transmission season in NYS, and chose representative, genetically distinct historic and contemporary virus strains to assess temperature-dependent vector competence in *Culex* mosquitoes. To do this, we amplified eight strains of WNV isolated in NYS, four from historic years with lower average temperatures (2003, 2004) and four from recent years with higher average temperatures (2017, 2018), and measured vector competence of colonized *Culex pipiens* at 20°C, 24°C, and 28°C, bracketing historic, contemporary, and future average summertime temperatures in NYS. These data were then utilized to calculate the relative R_0_ and estimate strain-specific thermal performance curves. Our results demonstrate temperature and strain-specific differences which are pertinent to WNV transmission models.

## Results

### Genetic Analysis

Genetic analysis of mean pairwise distance within transmission seasons, or annual amino acid diversity, was calculated using sequence data from WNV isolates acquired from NYS mosquito surveillance from 2000-2018[16,20]. When compared to the mean temperature during the transmission season (May – October), a positive correlation was identified between intraseason genetic diversity and temperature (Fig.1; r = 0.6, Pearson’s correlation, *p* < 0.05).

**Figure 1.**
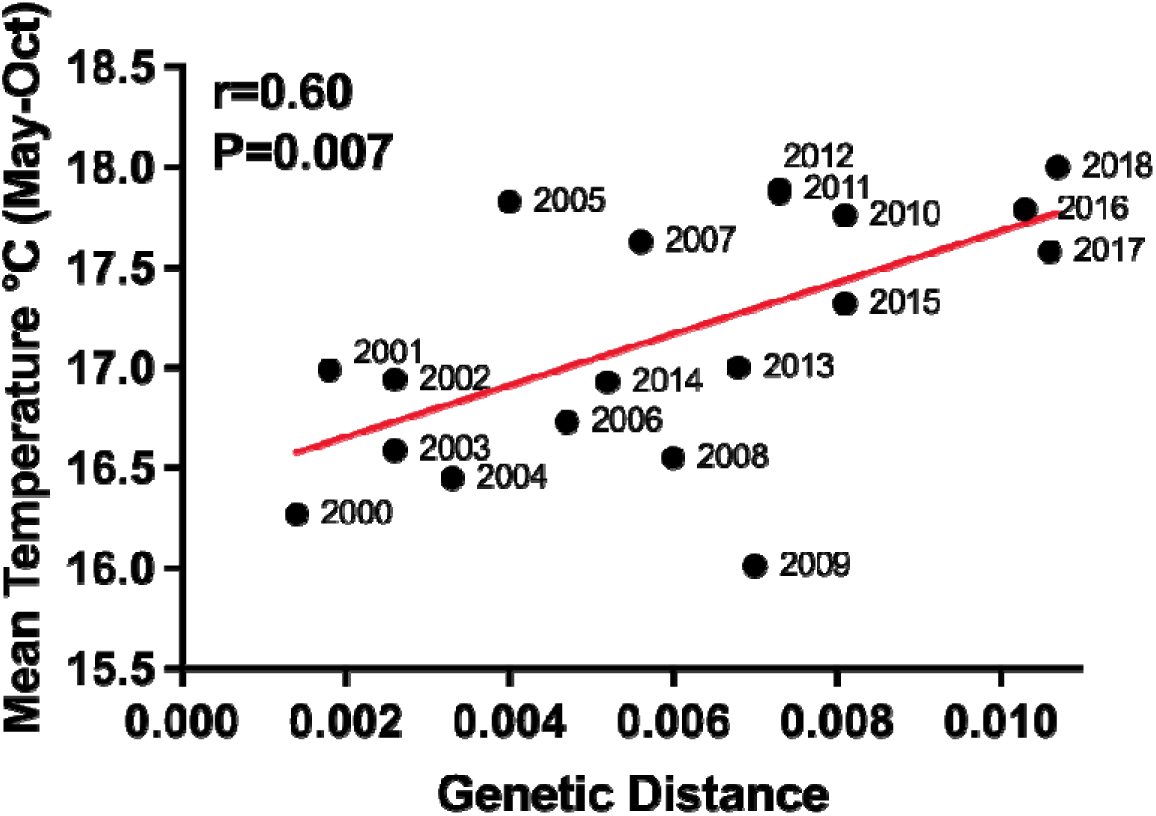
Genetic diversity of WNV strains in NYS. Correlation between genetic distance and mean temperature during transmission season. A total of 545 strains were sequenced and within-year amino acid distance was calculated in MEGA X. A positive correlation was identified between time and genetic distance (*p* < 0.05, Pearson correlation r = 0.60).

Pairwise amino acid differences among the eight experimental strains range from 1 – 6 amino acids (Fig. 2 and Table. 1). The majority of amino acid changes are in the non-structural proteins, particularly NS5. The four strains from 2003 and 2004 belong to the WN02 genotype, whereas the four strains from 2017 and 2018 share 2 amino acid changes among them, R1331K and I2513M, which are the substitutions defining the NY10 genotype[20].

**Figure 2.**
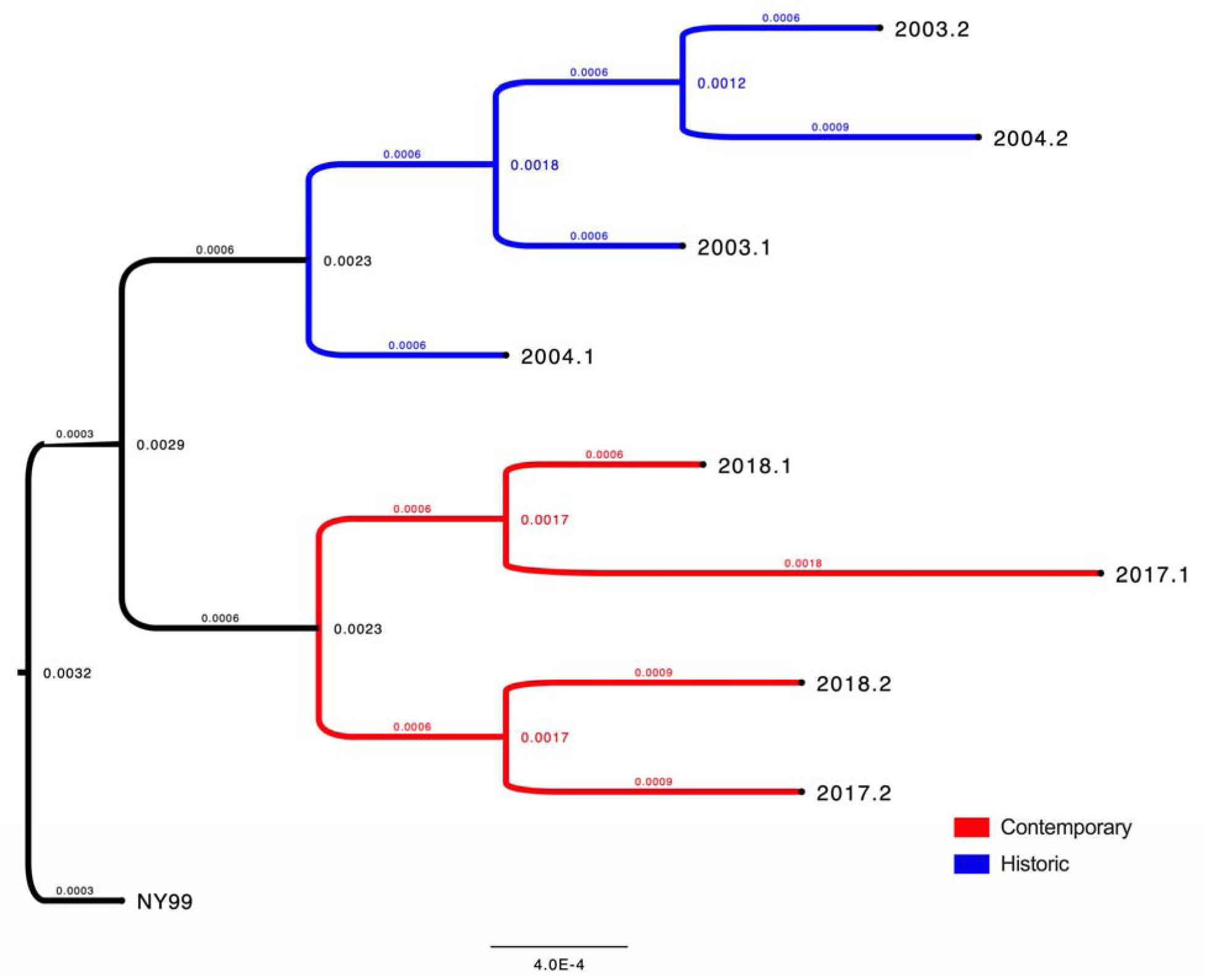
Phylogenetic tree of strains utilized in this study. A maximum likelihood phylogenetic tree of strains utilized in this study illustrates the genetic relationship on the nucleotide level. The root of this phylogenetic tree is the NY99 strain first identified in New York State in 1999, when West Nile virus was originally introduced in the Western hemisphere. Historic strains (2003-4) from cooler transmission seasons of 16°C are shown in blue and contemporary strains (2017-18) from warmer transmission seasons of 17°C are shown in red.

**Table 1.**
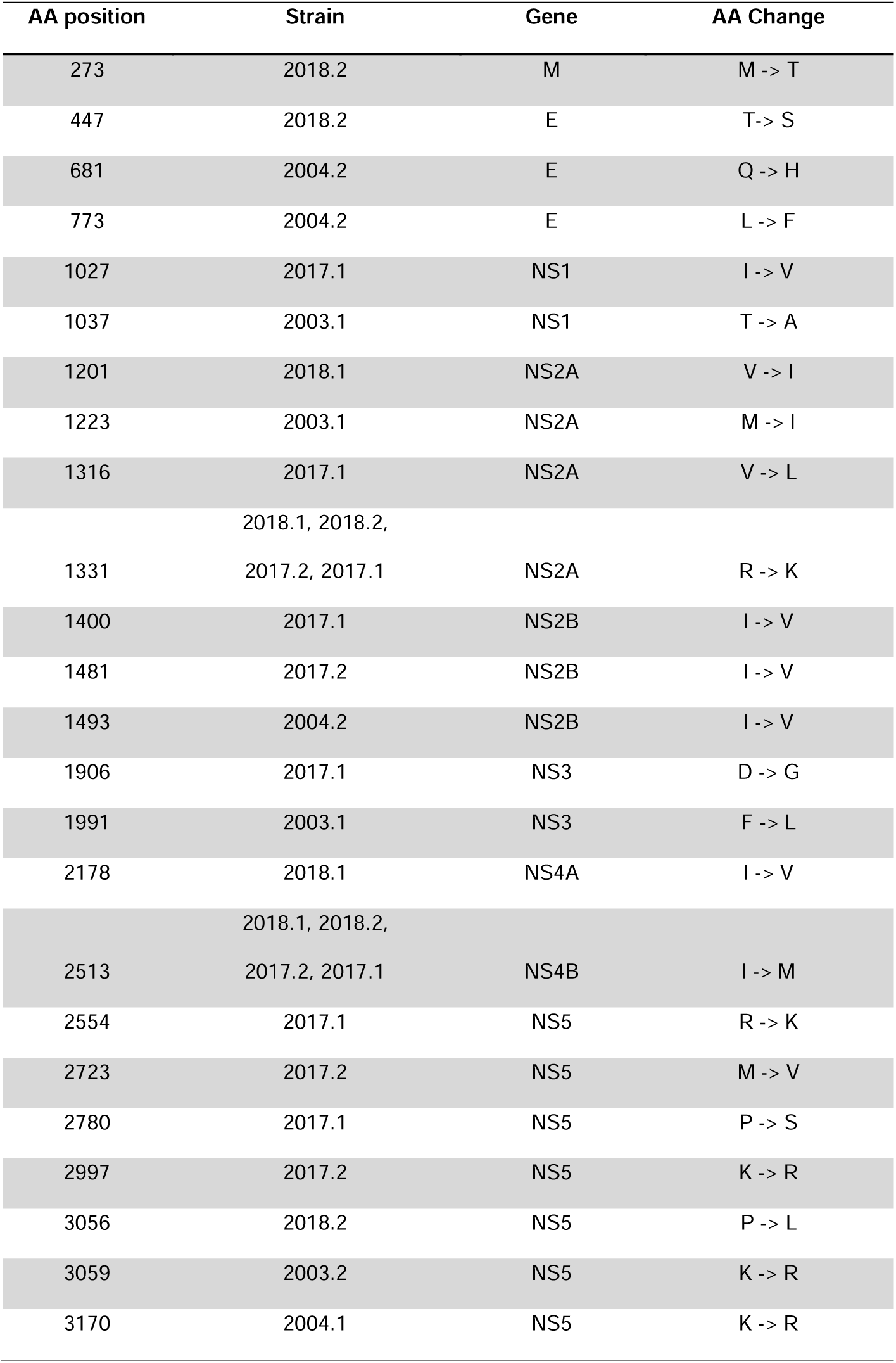
Amino acid changes identified in West Nile virus strains used in this study. This table depicts the amino acid position, strain designation, gene, and substitution identified

### Vector Competence

*Cx. pipiens* mosquitoes were fed on bloodmeals with 8 distinct strains of WNV and then housed at either 20°C, 24°C, or 28°C post feeding to determine if there is evidence for differential temperature-dependent vector competence for contemporary strains when compared to historic strains. These temperatures were chosen because they capture the realistic temperature range during transmission season (July-August) under historic, contemporary, and future predicted temperatures[21,22]. Mosquitoes were harvested on days 5 and 12 post-feeding (dpf) to determine the percentage of mosquitoes infected with the virus (indicated by positive bodies) or with disseminated infection (indicated by positive legs). These data show that *Cx. pipiens* vector competence is variable across temperatures and significant differences among strains exist (Fig.3 and Table.S1; *p* < 0.05, Chi-square test and 2-way ANOVA). Specifically, strain has a significant effect on infection rates at 5 dpf (*p* < 0.01) and 12 dpf (*p* <0.0001)(Table. 1; *p* < 0.05, 2-way ANOVA). At 20°C, *Cx. pipiens* had the highest dissemination rate for the strain 2004.1 at 5 dpf and 2017.1 at 12 dpf and the highest infection rates for the 2018.1 strain at 5 dpf and 12 dpf. At 24°C, *Cx. pipiens* had the highest dissemination rates for 2018.1 at 5 dpf and strain 2017.2 at 12 dpf, and the highest infection rate for 2018.1 at 5 dpf and 2004.2 at 12 dpf. At 28°C, *Cx. pipiens* had the highest dissemination rate for the 2018.1 strain at 5 dpf and 2004.2 strain at 12 dpf, and the highest infection rate for strain 2003.1 at 5 dpf and 2018.1 and 2003.1 at 12 dpf. When strain subsets (historic vs. contemporary) and days (5 vs. 12 dpf) were combined within temperatures, *Cx. pipiens* showed increased dissemination of contemporary strains at all temperatures (Fig.3 C; *p* < 0.05, Chi-square test and 2-way ANOVA), and temperature had a significant positive effect on dissemination (*p* < 0.01).

**Figure 3.**
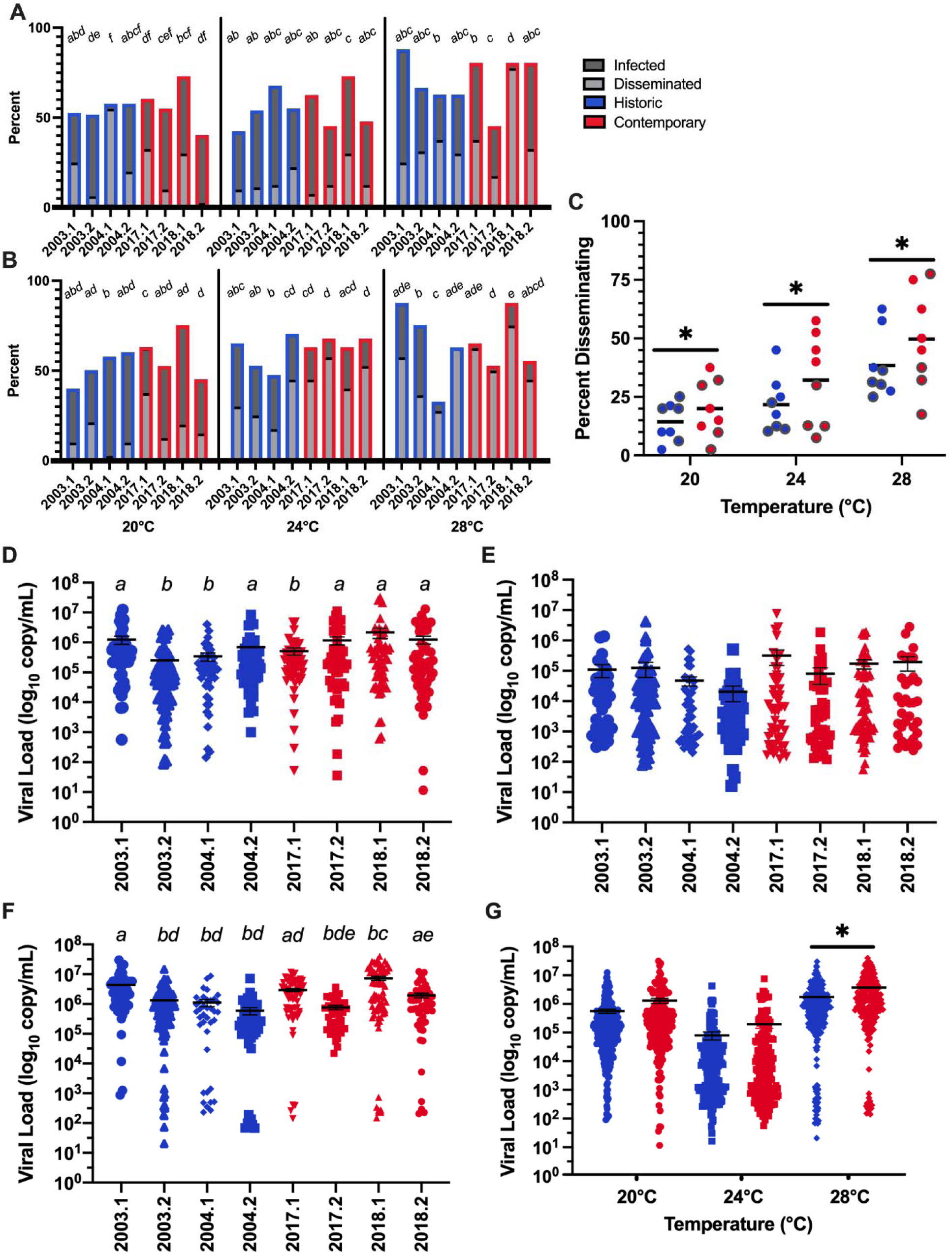
Vector competence and viral load measured in *Cx. pipiens* following blood feeding with historic and contemporary strains of West Nile virus from New York State at various temperatures. The percent of mosquitoes that were uninfected, infected, and infected with disseminating infection at **(A)** 5 days post-infection and (**B**) 12 days post-infection. Mosquitoes were fed 7.5 log_10_/mL WNV via blood meal using 8 previously identified WNV strains from NYS. Unique letters indicate statistically different measures (* *p* < 0.05, Chi-square test). (**C**) Percent of mosquitoes with disseminated infections within historic or contemporary strain subsets and combined time points. Gray outlined circles indicate 5 days post-infection and no outline indicates 12 days post-infection. Significant differences between total disseminating and total not disseminating mosquitoes were identified within temperature between strain subsets (* *p* < 0.05, Chi-square test). (**D**) Mosquitoes were held at 20°C, (**E**) 24°C, or (**F**) 28°C following infection. West Nile virus viral load (copy number equivalents) was determined in whole mosquitoes by using West Nile virus-specific quantitative reverse transcription PCR and standards. Viral load in individual mosquitoes after feeding (7.28-7.44 log_10_ pfu/mL) at both 5 and 12 days post-infection, as well as the mean +/-SEM are shown. Unique letters indicate statistically different measures between West Nile virus strains (\**p* < 0.05, one-way ANOVA with Šídák’s multiple comparisons test). (**G**) Combined viral load of *Cx. pipiens* exposed to WNV by day, temperature, and strain subsets. A significantly higher viral load was identified for combined contemporary strain data at 28°C (\**p* < 0.05, one-way ANOVA with Šídák’s multiple comparisons test).

Viral load in *Cx. pipiens* bodies were determined via qRT-PCR and significant differences among strains were identified at 20°C and 28°C (Fig.3 DF; *p* < 0.05 One-way ANOVA with Šídák’s multiple comparisons test). Additionally, a significant difference in mean viral load was found when strain subsets and time points were combined within temperature (Fig.3 G; *p* < 0.05, One-way ANOVA with Šídák’s multiple comparisons test). *Cx. pipiens* infected with contemporary WNV strains had higher viral loads compared to that of historic strains at 28°C. These results suggest that contemporary strains replicate more efficiently at higher temperatures.

### Thermal Performance Curves

Although estimating thermal performance curves with limited temperature data is challenging, we worked to refine previous estimates[23] of thermal performance curves (TPCs) for WNV dissemination in *Cx. pipiens.* To do this we used a Bayesian approach to identify strain-specific differences utilizing the vector competence data for each strain, using priors that conservatively assume no differences among strains in thermal performance and allowing the data presented here to determine any strain-specific differences. With these strain-specific TPCs, we obtained estimates of the temperature dependence of *Cx. pipiens* vector competence and R_0_ for contemporary and historic WNV strains at other temperatures than those directly measured (see Methods and Appendix for details). While the inferred vector competence TPCs had high uncertainty due to the limited temperature range measured for most strains (Fig. 1, Fig. S1, and Table S2), we found evidence for an overall increase in peak *Cx. pipiens* vector competence in the contemporary strains (Fig. S1), with a posterior probability of 0.955 that mean peak vector competence is higher in contemporary strains (probability of disseminated infection = 0.614; 95% credible interval:[0.443, 0.732]) compared to historic strains (0.437; 95%CI: [0.307, 0.583]). Disaggregating the strains, we also found increases in peak vector competence of various contemporary strains compared to historic strains (Fig. S2). Specifically, strains 2017.1, 2017.2, and 2018.1 vs. 2003.2, and strains 2017.1 and 2018.1 vs. 2004.1 have non-overlapping 95% CIs. Further, our results suggest that the mean vector competence thermal optima for the contemporary strains (30.8°C; 95%CI:[28.6°C, 33.2°C]) may be higher than for the historic strains (28.6°C; 95%CI:[26.5°C, 31.0°C]), although this result is inconclusive (posterior probability of 0.912) and needs to be confirmed in future studies with measurements at a wider temperature range. Lastly, we calculated temperature-dependent relative R_0_ by applying the new vector competence thermal performance curves into a previously published model by Shocket *et al* 2020[23] (see Appendix). We did not find clear differences in the thermal optimum or thermal limits for transmission between historic and contemporary WNV strains (Fig. S3 and Table. S3). When we compare the relative R_0_ across temperatures, we found increased transmission of contemporary strains relative to historic strains at high temperatures, starting at around 23°C (Fig. S4).

### Temperature Specific Transmission

To assess transmission potential at each of the temperatures for which we measured vector competence, we calculated the relative R_0_ using previously published temperature-specific *Cx. pipiens* life history trait estimates^23^ along with the new strain-specific vector competence data (Table.S1). We found that, on average, contemporary WNV strains have higher transmission at 24°C and 28°C compared to the historic strains (Fig.4 AB). To compare strain subsets, we calculated the ratio of the mean relative R_0_ of both subsets and found contemporary strains have a 29.3% 95%CI:[12.8%, 48.0%] increase in transmission potential at 24°C (with a >0.999 posterior probability of higher transmission in contemporary strains) and a 10.2% 95%CI:[−0.3%, 21.5%] increase at 28°C (with a 0.971 posterior probability that transmission is higher in contemporary strains). Although there is variability among strains (Fig. S5), these data demonstrate that, on average, *Cx. pipiens* have increased transmission potential for contemporary WNV strains compared to historic strains at the higher temperatures evaluated.

**Figure 4.**
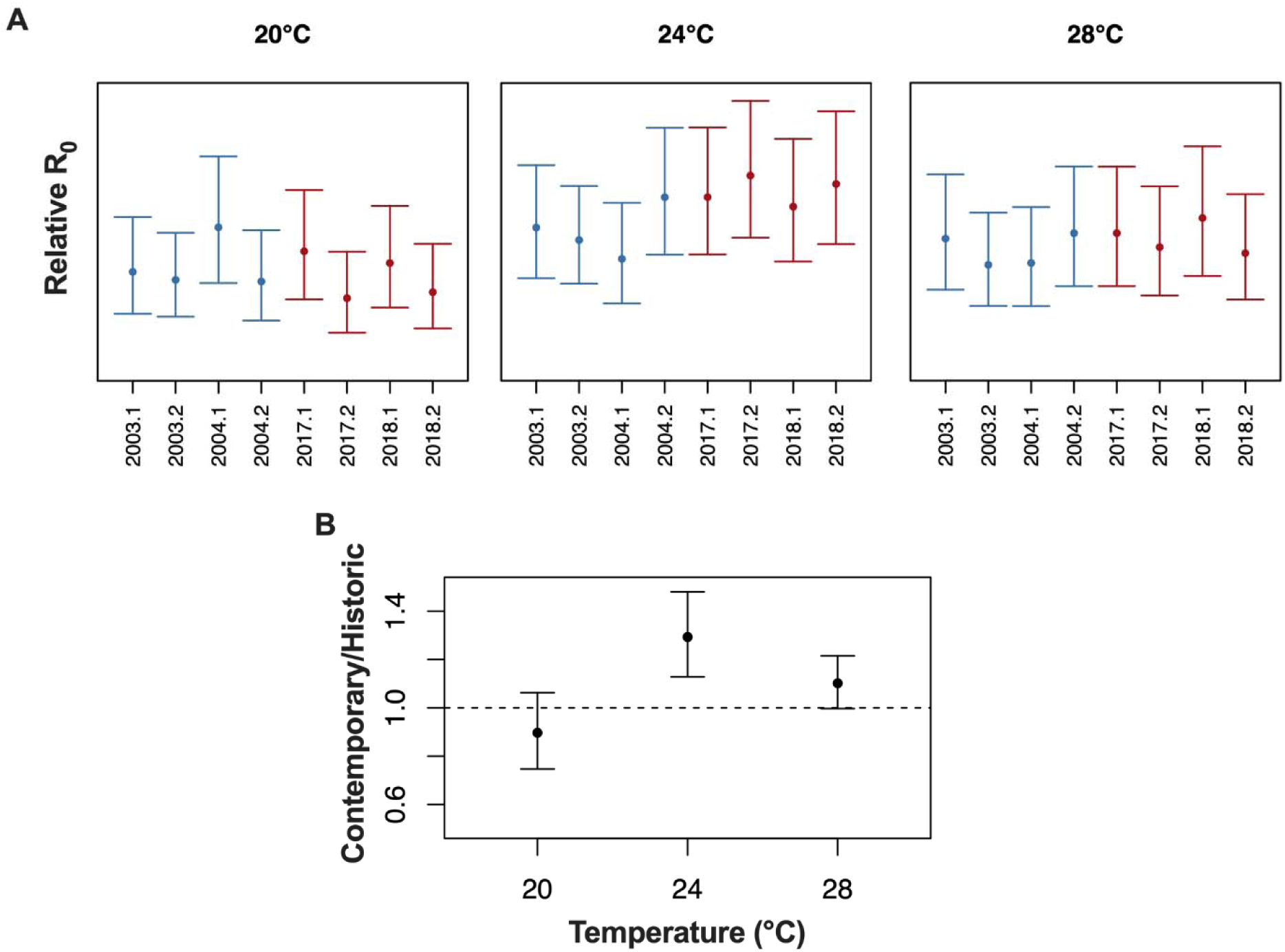
Point-wise relative R_0_ of historic and contemporary West Nile virus strains in *Cx. pipiens*. (**A**) Relative R_0_ for historic (blue) and contemporary (red) WNV strains in *Cx. pipiens* is shown at 20°C, 24°C, and 28°C. Posterior means are shown as points and 95% credible intervals as lines. (**B**) The ratio between the mean relative R_0_ of the four contemporary strains and the mean of the four historic strains is shown at each temperature. Points correspond to posterior means and lines to 95% credible intervals. A ratio of one (corresponding to equal transmission of contemporary and historic strains) is shown as a dotted line.

## Discussion

The data presented here demonstrate significant strain-specific variability in temperature-dependent WNV transmissibility. Analysis of genetic data suggests temperature-associated diversification of WNV in New York State (Fig. 1), and previous studies suggest that this diversification was to some extent driven by adaptive evolution[16,20]. To test the hypothesis that increased temperatures contributed to this adaptation we retrospectively assessed WNV thermal sensitivity using eight genetically distinct WNV strains from the years 2003-2004 and 2017-2018 (Fig. 2). *Cx. pipiens* have increased vector competence for contemporary WNV strains across temperatures (Fig. 3C and Table.S1). Data from this experiment and mosquito life history traits from previously published work[24] were then used to calculate the relative R_0_ point-wise at 20°C, 24°C, and 28°C. *Cx. pipiens* have increased transmission potential for contemporary WNV strains compared to historic strains at 24°C and 28°C (Fig. 4B). Overall, WNV populations that evolved more recently and under increased temperature demonstrate increased genotypic and phenotypic diversity. Furthermore, our data suggest that as temperature increases, strains with fitness advantages at higher temperatures will continue to become more dominant and drive increased transmission by *Cx. pipiens*.

Temperature was previously identified as a primary driver of genetic divergence of WNV on a national scale[17], although numerous selective and stochastic factors influence diversity. Temperature is known to increase viral replication of an already error-prone virus, thereby increasing the mutation frequency[18]. Host distribution and abundance, be it avian or arthropod, may also influence genetic diversity, and these factors will also shift with temperature variations (though in potentially idiosyncratic ways)[25,26]. WNV is currently endemic in NYS and many different strains circulate simultaneously on regional and local scales[20,27]. This study utilized a representative group of eight WNV strains, chosen due to temporal separation, but also due to differences in seasonal temperature and WNV prevalence. While these strains are not an exhaustive representation of WNV diversity, they represent the dominant genotypes circulating at the time. The majority of the amino acid changes identified among strains were found in the non-structural proteins, suggesting a role in immune evasion, viral replication, and/or assembly[28–30]. Mutations in the envelope (E) protein have been shown to regulate conformational changes pertinent to WNV infectivity in mammalian cell lines[31]. Studies of NS1 mutations identified their role in viral replication and secretion[32]. The NS2 protein is involved in the biogenesis of virus-induced membranes, suggesting an important role in virus assembly[29], as well as RNAi suppression [33]. Substitutions in the NS3, which encodes the viral helicase and ATPase, have been shown to influence transmission by mosquitoes[34]. The NS4 protein interacts with several host and viral proteins, and mutations within the protein can have significant effects on host-specific fitness and immunity[35]. The NS5 gene, which encodes the viral polymerase and methyltransferase, controls viral replication and mutation rate. Substitutions in these proteins have been shown to influence host-specific viral fitness[28]. More work is necessary to elucidate the role of specific mutations identified in this study and how temperature might influence phenotypic consequences.

As previously shown, mosquito vector competence has a non-linear relationship that increases until approximately 26-30°C and decreases above those temperatures[23]. Despite this, genotype and strain-specific differences have been previously identified in the context of WNV transmission[8,15,16,36–38]. Although we demonstrate that, on average, *Cx. pipiens* has increased vector competence for contemporary strains, but differences were not uniform within historic and contemporary strain subsets. For instance, the strain with the highest infection rate was historic (2003.1), although increased infection only occurred at 28°C. The strain with the highest dissemination rate at 28°C was 2018.1, which also showed increased competence at 20°C and 24°C. Previous *in vitro* studies predicted that evolution at higher temperatures should result in more variability and broad adaptation[18]. This is consistent with the results presented here. These data demonstrate that strain-specific differences may have important impacts on WNV transmission, particularly in the context of climate warming. Additional consideration should be given to the mosquito population and the potential for unique genotype-genotype-temperature interactions[39–41].

The typical pattern of arbovirus infection in mosquitoes is for dissemination rates to increase at later time points as the virus has more opportunity to replicate and escape the midgut epithelial barrier[16]. However, in this study, we measured decreased dissemination at 12 dpf compared to 5 dpf for some of the strains utilized in this study. We hypothesize that lower measured dissemination rates at 12 dpf may be due to the death of infected mosquitoes, especially at higher temperatures. Mosquitoes with high viral loads may have died earlier in the study, as has been noted in previous studies with WNV in *Cx. pipiens*[38], resulting in a lower measured fraction of disseminated infection in the remaining mosquito pool. Accounting for mosquito death upon infection may be pertinent to future experimental protocols and to models aiming to capture the impact of high temperature on transmission. To avoid underestimating dissemination in our R0 models in these cases, we used the highest level of dissemination recorded for each treatment, which occurred at either 5 or 12 dpf, as our estimate of dissemination probability.

In this study, we used a combination of novel empirical and previously published data to calculate the transmission potential of 8 WNV strains. The calculation of relative R_0_ is a direct estimate of transmissibility but does not consider vertebrate host abundance and recovery rate. Importantly, a previous study found that WNV NY10 strains genetically similar to the contemporary strains used here sustained higher levels of viremia in American robins[16]. These data suggest that the strain-specific transmission advantage may be more pronounced for contemporary strains than is estimated from vector traits alone[23].

Vector competence was assessed at three different temperatures: 20°C, 24°C, and 28°C, which span the average temperature range during peak transmission season from the isolation year of each strain as well as a predicted future temperature (28°C) based on climate modeling[22]. This temperature range was reflective of means during peak transmission, but higher and lower extremes may be of interest given daily and seasonal fluctuations[23]. Alterations to transmissibility at lower temperatures, in particular, could have a significant impact on the duration of the WNV transmission season in temperate regions such as NYS. Moreover, given the temperature range measured in this study, there is substantial uncertainty in the inferred strain-specific TPCs for dissemination, especially at high temperatures. Because we did not have empirical measurements above 28°C, the estimated thermal optima and maxima from TPCs for dissemination are largely influenced by priors based on previous TPC fits for other WNV strains and may underestimate the true variation across the strains studied here[23]. Additionally, cycling temperatures more accurately mimicking daily fluctuations have been shown to yield differing results compared to static temperature for transmission of WNV and other flaviviruses[42,43]. As WNV in nature continues to evolve under increased and varying temperatures, mosquitoes in nature will experience such pressures[44]. Future studies should investigate *Cx. pipiens* ability to adapt to increasing temperature and how these selective pressures might impact WNV transmission.

The risk of WNV is predicted to increase in some regions as temperature increases[45] yet predictive models that do not account for the potential for increased transmissibility of emerging WNV strains may underestimate the future risk. Refining WNV thermal biology to consider the diverse and evolving nature of vector-virus interactions more accurately is key to preparing public health efforts for future vector-borne disease risks.

## Methods

### West Nile Virus Mosquito Surveillance

Mosquitoes were collected in Centers for Disease Control (CDC) light traps by NYS county health departments and speciated pools were submitted to the NYS Arbovirus Laboratory for processing and testing. Pools consisted of 15 - 60 *Cx. pipiens* and/or *Cx. restuans* females in 1 mL mosquito diluent [20% heat-inactivated fetal bovine serum (FBS) in Dulbecco’s phosphate-buffered saline (PBS) plus 50 μg/mL penicillin/streptomycin, 50 μg/mL gentamicin, and 2.5 μg/mL Fungizone] with 1 steel bead (Daisy Outdoor Products, Rogers, AR). Pools were processed by homogenization for 30 seconds at 24 Hz in a Mixer Mill MM301 (Retsch, Newtown, PA), followed by centrifugation at 6000 rcf for 5 minutes. WNV-positive pools were identified by quantitative real-time reverse transcription polymerase chain reaction (qRT-PCR)[46].

### Sequencing and genetic analyses

Sequencing was performed on the Illumina MiSeq platform (San Diego, CA) following the Bialosuknia et al. protocol[20]. Paired-end reads were assembled to a WN02 genotype reference (DQ164190) deploying Geneious Pro’s reference mapping tool using high sensitivity and free end gaps with 10 iterations of fine-tuning, trimming paired read overhangs. The same parameters were used to map reads to the consensus assembly. Alignment was completed with Geneious Pro, with the algorithm set to the slow and accurate L-INS-I alignment algorithm, with the scoring matrix set to 200PAM/K=2. The gap open penalty was set to 1.53, and the offset value was set to 0.123. Annual WNV amino acid diversity was calculated using the software Molecular Evolutionary Genetics Analysis (MEGA X) by computing the within-group mean genetic distance. Daily mean temperatures were derived from GridMET[21], using the GridMET downloader tool[22] and averaged over the entire period. A maximum likelihood phylogenetic tree of strains utilized in this study was generated in Geneious Pro and edited in Fig Tree v1.4.4. The root of this phylogenetic tree is the NY99 (AF196835.2) strain first identified in New York State in 1999.

### Viruses

The WN02 strain used (2003.2) was isolated in 2003 (DQ164189) from an American crow (*Corvus brachyrhynchos*) found in Albany County, NY which was initially amplified on Vero cells for sequencing, and then later amplified on C6/36 cells (*Aedes albopictus*, ATCC, Manassas, VA) for downstream use. All other strains utilized in this study were isolated from *Culex* mosquito surveillance pools. The samples were collected in several counties across NYS; 2004.1 (KX547454), 2003.1 (KX547310), 2003.2 (DQ164189), 2004.2 (KX547408.1), 2018.1 (MT968016.1), 2017.1 (MT968008.1), 2018.2 (MT968014.1), and 2017.2 (MT967993.1). Strains were either amplified one time on Vero and C6/36 cells or two times on C6/36 only. After 5 days of amplification on C6/36 tissue culture supernatant was harvested and stored in 20% FBS at −80°C.

### Mosquito Rearing

*Cx. pipiens* colony mosquitoes were originally collected in Pennsylvania in 2004 (courtesy of M. Hutchinson) and have been maintained in the colony at the Wadsworth Center Arbovirus laboratory. *Cx. pipiens* mosquitoes were maintained in 30.5-cm^3^ cages in an environmental chamber at 27 ± 2°C with a relative humidity of 45-65% and a photoperiod of 16:8 (L:D) h and provided cotton pads with 10% sucrose *ad libitum*.

### Vector Competence

To assess WNV strain-specific competence, an estimated 5000 four-to-seven-day-old adult females were collected and fed on doses of WNV ranging from 7.28-7.44 log_10_ pfu/mL. Bloodmeals consisted of a 4:1 mixture of diluted virus stock and chicken blood (Colorado Serum Company, Denver, CO), and a final concentration of 2.5% sucrose. Following one hour of feeding using an artificial feeding chamber (Hemotek, Blackburn, UK) at 37°C, mosquitoes were anesthetized, and the engorged females were collected and held at 20°C, 24°C, or 28°C. These temperatures were shown to mimic regional temperatures relevant to these unique populations and a predicted future temperature. A vector competence assay was performed at 5- and 12-days post-feeding (dpf). Individual mosquitos’ legs and body were saved separately with a 4.5 mm zinc-plated steel ball (Daisy, Dallas, TX) in 500 uL mosquito diluent (PBS w/ 20% FBS, 100 μg/mL penicillin/streptomycin, 10 μg/mL gentamicin, 1 μg/mL amphotericin B) at −80°C. This experiment was performed in 2 different groups with the WN02 strain serving as the positive control in each group. To determine positivity, thawed samples were homogenized at 24Hz for 3 minutes, followed by centrifugation at 1200 rpm for 3 minutes and RNA was extracted using a MagMAX viral isolation kit (Applied Biosystems, Waltham, MA, USA) on a Tecan Evo 150 liquid handler (Tecan, Mannedorf, Switzerland). Real-time quantification RT-PCR was completed using TaqMan One-Step RT-PCR master mix (Applied Biosystems, Waltham, MA, USA) and analyzed on Quant Studio 5 (Thermo Fisher, Waltham, MA, USA). WNV primers and probes were designed as previously described and copy standards were utilized for quantification [47]. A total of 40 mosquitoes were tested for each strain at each timepoint for a total of 2160 tested mosquitoes. Vector competence was assessed by quantifying the proportions of infected (positive bodies) and disseminated (positive legs) at each time point. Strain-specific data were analyzed and compared using GraphPad Prism 9.

### Thermal Limits Experiments

Additional data points from another experiment were used to better estimate the thermal limits of WNV transmission (see Appendix for data). Methods for such experiments are as follows: the 2003.1 and 2017.1 strains were grown as described above. *Cx. pipiens* colony mosquitoes were originally collected in Pennsylvania in 2004 (courtesy of M. Hutchinson) and have been highly colonized at the Wadsworth Center Arbovirus laboratory and were reared as outlined above.

Four to seven-day-old adult females were collected and fed on doses of WNV ranging from the 2003.1 strain 6.92 Log_10_ pfu/ml or 2017.1 strain 7.38 Log_10_ pfu/ml. Blood meals consisted of a 4:1 mixture of diluted virus stock and chicken blood (Colorado Serum Company, Denver, CO), and a final concentration of 2.5% sucrose. Additionally, a non-infectious blood meal was also provided to separate mosquitoes to allow the comparison of WNV-exposed and unexposed life history traits. Following one hour of feeding using an artificial feeding chamber (Hemotek, Blackburn, UK) at 37°C, mosquitoes were anesthetized, and the engorged females were collected, knocked down using CO_2,_ and placed in individual 50 mL conical tubes in Styrofoam rack, with a hole in the top of the conical covered in mesh, a hole in the bottom of the conical to allow addition of water for laying, and a dental dam around bottom of the conical to hold the tube in place. Cotton pads soaked in 10% sucrose ad libitum were placed on top of the conicals. Individual conicals were then held at constant temperatures of 15°C and 33°C. 45-50 blood-fed mosquitoes for each WNV strain and temperature were utilized.

Survival was checked daily for all treatment and temperature groups. The bottom of conicals were checked daily for eggs; when eggs were observed in a conical, the egg raft was removed using a wooden stick. Additional water was added as needed to the bottom of the conicals. Mosquitoes were starved overnight for 12-24 h and offered 200-ul defibrinated chicken blood (Colorado Serum Company, Denver, CO) with 2.5% sucrose via absorbent pad for 2-h 1x per week. Mosquitoes were monitored for feeding activity after the 2 hours by eye.

Upon death, individual mosquito’s legs and body were saved separately with a 4.5 mm zinc-plated steel ball (Daisy, Dallas, TX) in 500 μL mosquito diluent (PBS w/ 20% FBS, 100 μg/mL penicillin/streptomycin, 10 μg/mL gentamicin, 1 μg/mL amphotericin B) at −80°C. To determine positivity, thawed samples were homogenized at 24Hz for 3 minutes, followed by centrifugation at 1200 rpm for 3 minutes, and RNA was extracted using a MagMAX viral isolation kit (Applied Biosystems, Waltham, MA, USA) on a Tecan Evo 150 liquid handler (Tecan, Mannedorf, Switzerland). Real-time quantification quantitation RT-PCR was completed using TaqMan One-Step RT-PCR master mix (Applied Biosystems, Waltham, MA, USA) and analyzed on Quant Studio 5 (Thermo Fisher, Waltham, MA, USA). WNV primers and probes were designed as previously described and copy standards were utilized for quantification [47]. 45-50 mosquitoes were tested for each genotype. Vector competence was assessed by quantifying the proportions of infected (positive bodies) and disseminated (positive legs). Genotype-specific data were analyzed and compared using GraphPad Prism 9.

### Relative R_0_ Estimates

Following previous work, we evaluate the effects of strain-specific vector competence in WNV transmission by combining various mosquito and WNV life history traits to estimate a temperature-dependent relative R_0_ with a Bayesian approach[23]. Relative R_0_ is defined as

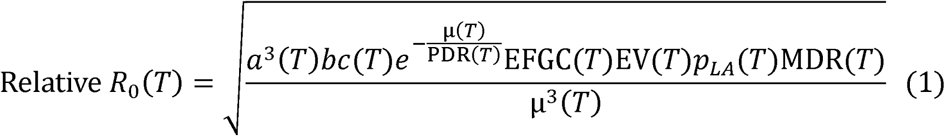

where *a*(*T*) is biting rate, *bc*(*T*) vector competence, μ(*T*) the mosquito death rate, PDR(*T*) the pathogen development rate, EFGC(*T*) eggs per female per gonotrophic cycle, EV(*T*) egg viability, and *p_LA_*(*T*) the proportion of mosquito larvae that survive to adulthood. Our approach is to combine strain-specific vector competence estimates (from this work) with previously measured *Cx pipiens* life history traits[23].

Estimating a thermal performance curve for vector competence with limited temperature treatments is challenging and requires additional assumptions in the form of strongly informative prior distributions. Because of this, we conducted two analyses to estimate relative R_0_. In the main text, we present a point-wise analysis where relative R_0_ was estimated only at the three temperatures for which vector competence was measured experimentally for all viral strains, which does not require estimating a TPC for vector competence but limits conclusions to the specific temperatures measured directly.

An alternate analysis where we combined strong priors informed by previous experiments and a hierarchical model with partial pooling of data of multiple strains to estimate strain-specific TPCs for vector competence is presented in the Appendix. In this alternate analysis, we combined the estimated vector competence TPCs with previously estimated mosquito life history traits to estimate relative R_0_ curves as a function of temperature. For both analyses, the maximum dissemination proportion of the two measured time points (5 or 12 dpf) was used for the vector competence measurement at each temperature.

### Point-wise Vector Competence and Relative R_0_ Estimates

Vector competence was estimated separately for each strain at 20°C, 24°C and 28°C. The number of mosquitoes with a disseminated infection was modeled with a binomial likelihood, with a uniform prior for the disseminated proportion at each temperature (see Appendix). Strain-specific relative R_0_ estimates were obtained by combining the vector competence estimates with previously estimated mosquito life history traits and WNV development rate[23].

### Vector Competence TPCs and Relative R_0_ Curves

In an alternate analysis, a Bayesian approach was used to estimate thermal performance curves (TPCs) for vector competence for the WNV strains described in this work. Additional points at thermal extremes of 15°C and 33°C were added for two strains from other experiments, specifically 2003.2 and 2017.1 (see *Thermal Limits Experiments* section in Methods and additional data in Appendix). As in previous work that fit TPCs to dissemination proportions for WNV in *Cx. pipiens* and other *Culex* mosquitoes[23], we used a quadratic model as the functional form of the TPC model. A hierarchical model was used to estimate the TPC parameters of each strain. This performs partial pooling using the data from multiple strains by assuming that the TPC parameters for each individual strain come from a common probability distribution for either historic or contemporary strains (see Appendix for details). Informative prior distributions based on previous TPC fits to dissemination proportion in *Cx. pipiens*[23] were used to constrain the thermal limits and maximum dissemination proportion to reasonable values. Details regarding the statistical model, including all prior distributions and how they relate to previous estimates, can be found in the Appendix.

Models were fit using Markov Chain Monte Carlo (MCMC) with the r2jags R package[48], an interface for the JAGS (Just Another Gibbs Sampler) program[49]. Four independent MCMC chains were run for 200,000 iterations, discarding the first 40,000 iterations as burn-in[50]. The resulting MCMC chains were thinned, saving every eight iterations. Chain convergence was monitored both by visual inspection of trace plots factor *R̂* < 1.01 for all parameters. A summary of the posterior distribution (mean, 95% and density plots of the individual chains and by ensuring the potential scale reduction credible interval [CI], *R̂*, *n_eff_*) for all model parameters corresponding to the TPC fit for each trait is available in the Appendix. Posterior distributions for optimal temperatures were calculated analytically using the posterior samples of the model parameters for each strain (see Appendix). The consequences of the strain-specific differences in vector competence on the temperature dependence of the basic reproductive number R_0_ for WNV for each historical and contemporary virus strain were estimated with the relative R_0_ approach, as in Shocket *et al.* 2020[23] based on the thermal performance curves fit here and previously published estimates for the thermal performance of *Cx pipiens* life history traits and WNV pathogen development rate (see Appendix for details).

## Supplemental Figures and Tables

**Table S1.** *Cx. pipiens* vector competence for 8 representative West Nile virus isolates from New York State at 5 and 12 days after infectious bloodmeal at 20°C, 24°C, and 28°C.

| Temperature<br>(°C) | Strain | 5 dpf |  |  |  | 12 dpf |  |  |  |
| --- | --- | --- | --- | --- | --- | --- | --- | --- | --- |
|  |  | Infection<br>n | (%) | Dissemination<br>n | (%) | Infection<br>n | (%) | Dissemination<br>n | (%) |
| **22** | 2003.1 | 21 | 52.50% | 10 | 47.62% | 16 | 40.00% | 4 | 25.00% |
|  | 2003.2 | 41 | 51.25% | 5 | 12.20% | 40 | 50.00% | 17 | 42.50% |
|  | 2004.1 | 23 | 57.50% | 22 | 95.65% | 23 | 57.50% | 1 | 4.35% |
|  | 2004.2 | 23 | 57.50% | 8 | 34.78% | 24 | 60.00% | 4 | 16.67% |
|  | 2017.1 | 24 | 60.00% | 13 | 54.17% | 25 | 62.50% | 15 | 60.00% |
|  | 2017.2 | 22 | 55.00% | 4 | 18.18% | 21 | 52.50% | 5 | 23.81% |
|  | 2018.1 | 29 | 72.50% | 12 | 41.38% | 30 | 75.00% | 8 | 26.67% |
|  | 2018.2 | 16 | 40.00% | 1 | 6.25% | 18 | 45.00% | 6 | 33.33% |
| **24** | 2003.1 | 17 | 42.50% | 4 | 23.53% | 26 | 65.00% | 12 | 46.15% |
|  | 2003.2 | 43 | 53.75% | 9 | 20.93% | 42 | 52.50% | 20 | 47.62% |
|  | 2004.1 | 27 | 67.50% | 5 | 18.52% | 19 | 47.50% | 7 | 36.84% |
|  | 2004.2 | 22 | 55.00% | 9 | 40.91% | 28 | 70.00% | 18 | 64.29% |
|  | 2017.1 | 25 | 62.50% | 3 | 12.00% | 25 | 62.50% | 18 | 72.00% |
|  | 2017.2 | 18 | 45.00% | 5 | 27.78% | 27 | 67.50% | 23 | 85.19% |
|  | 2018.1 | 29 | 72.50% | 12 | 41.38% | 25 | 62.50% | 16 | 64.00% |
|  | 2018.2 | 19 | 47.50% | 5 | 26.32% | 27 | 67.50% | 21 | 77.78% |
| **28** | 2003.1 | 35 | 87.50% | 10 | 28.57% | 35 | 87.50% | 23 | 65.71% |
|  | 2003.2 | 53 | 66.25% | 25 | 47.17% | 60 | 75.00% | 29 | 48.33% |
|  | 2004.1 | 25 | 62.50% | 15 | 60.00% | 13 | 32.50% | 11 | 84.62% |
|  | 2004.2 | 25 | 62.50% | 12 | 48.00% | 25 | 62.50% | 25 | 100.00% |
|  | 2017.1 | 32 | 80.00% | 15 | 46.88% | 26 | 65.00% | 25 | 96.15% |
|  | 2017.2 | 18 | 45.00% | 7 | 38.89% | 21 | 52.50% | 20 | 95.24% |
|  | 2018.1 | 32 | 80.00% | 31 | 96.88% | 35 | 87.50% | 30 | 85.71% |
|  | 2018.2 | 32 | 80.00% | 13 | 40.63% | 22 | 55.00% | 18 | 81.82% |

**Table S2.** Estimated thermal limits and optima for vector competence of contemporary and historic WNV strains. For each strain, we report the posterior mean and 2.5% and 97.5% quantiles, representing a 95% credible interval of the minimum (Tmin), maximum (Tmax), and optimum/peak transmission (Topt) temperatures for strain-specific vector competence.

| Strain | Parameter | Mean | 2.5% | 97.5% |
| --- | --- | --- | --- | --- |
| **2003.1** | Tmin | 15.0 | 11.5 | 17.9 |
|  | Topt | 29.2 | 26.2 | 32.4 |
|  | Tmax | 43.3 | 37.8 | 45.1 |
|  | bc_max | 0.465 | 0.361 | 0.599 |
| **2003.2** | Tmin | 13.6 | 11.8 | 14.6 |
|  | Topt | 29.0 | 26.7 | 32.4 |
|  | Tmax | 44.5 | 39.7 | 51.3 |
|  | bc_max | 0.362 | 0.297 | 0.431 |
| **2004.1** | Tmin | 11.8 | 5.6 | 15.1 |
|  | Topt | 26.8 | 22.4 | 30.5 |
|  | Tmax | 41.8 | 33.8 | 48.2 |
|  | bc_max | 0.407 | 0.319 | 0.497 |
| **2004.2** | Tmin | 15.7 | 12.3 | 18.5 |
|  | Topt | 29.4 | 26.5 | 32.5 |
|  | Tmax | 42.7 | 38.2 | 47.3 |
|  | bc_max | 0.530 | 0.397 | 0.693 |
| **2017.1** | Tmin | 14.3 | 13.3 | 14.8 |
|  | Topt | 30.7 | 28.4 | 34.0 |
|  | Tmax | 47.2 | 42.5 | 54.0 |
|  | bc_max | 0.672 | 0.587 | 0.765 |
| **2017.2** | Tmin | 17.4 | 14.6 | 19.2 |
|  | Topt | 30.9 | 26.8 | 34.2 |
|  | Tmax | 44.4 | 35.3 | 51.3 |
|  | bc_max | 0.608 | 0.472 | 0.734 |
| **2018.1** | Tmin | 16.4 | 14.0 | 18.3 |
|  | Topt | 30.8 | 27.9 | 33.9 |
|  | Tmax | 45.1 | 39.1 | 51.5 |
|  | bc_max | 0.674 | 0.557 | 0.822 |
| **2018.2** | Tmin | 17.0 | 14.3 | 19.0 |
|  | Topt | 30.9 | 26.9 | 34.4 |
|  | Tmax | 44.7 | 36.2 | 47.0 |
|  | bc_max | 0.576 | 0.430 | 0.707 |

**Table S3.** Estimated thermal limits and optima for the relative R_0_ of contemporary and historic WNV strains. For each strain, we report the posterior mean and 2.5% and 97.5% quantiles, representing a 95% credible interval of the minimum (Tmin), maximum (Tmax), and optimum/peak transmission (Topt) temperatures for strain-specific relative R_0_.

| Strain | Tmin | 2.5% | 97.5% | Topt | 2.5% | 97.5% | Tmax | 2.5% | 97.5% |
| --- | --- | --- | --- | --- | --- | --- | --- | --- | --- |
| 2003.1 | 15.1 | 12.1 | 17.9 | 24.5 | 23.4 | 25.6 | 35.0 | 32.9 | 37.7 |
| 2003.2 | 13.8 | 12.2 | 15.1 | 24.3 | 23.3 | 25.5 | 35.0 | 32.9 | 37.7 |
| 2004.1 | 12.9 | 9.9 | 15.5 | 24.1 | 23.0 | 25.3 | 34.9 | 32.7 | 37.6 |
| 2004.2 | 15.7 | 12.6 | 18.5 | 24.6 | 23.5 | 25.7 | 35.0 | 32.9 | 37.6 |
| 2017.2 | 17.3 | 14.6 | 19.2 | 24.8 | 23.8 | 25.9 | 34.9 | 32.8 | 37.6 |
| 2017.1 | 14.3 | 13.4 | 15.1 | 24.5 | 23.5 | 25.6 | 35.0 | 32.9 | 37.7 |
| 2018.2 | 17.0 | 14.3 | 19.0 | 24.8 | 23.8 | 25.9 | 34.9 | 32.8 | 37.6 |
| 2018.1 | 16.4 | 14.0 | 18.3 | 24.7 | 23.7 | 25.8 | 35.0 | 32.9 | 37.7 |

**Figure S1.**
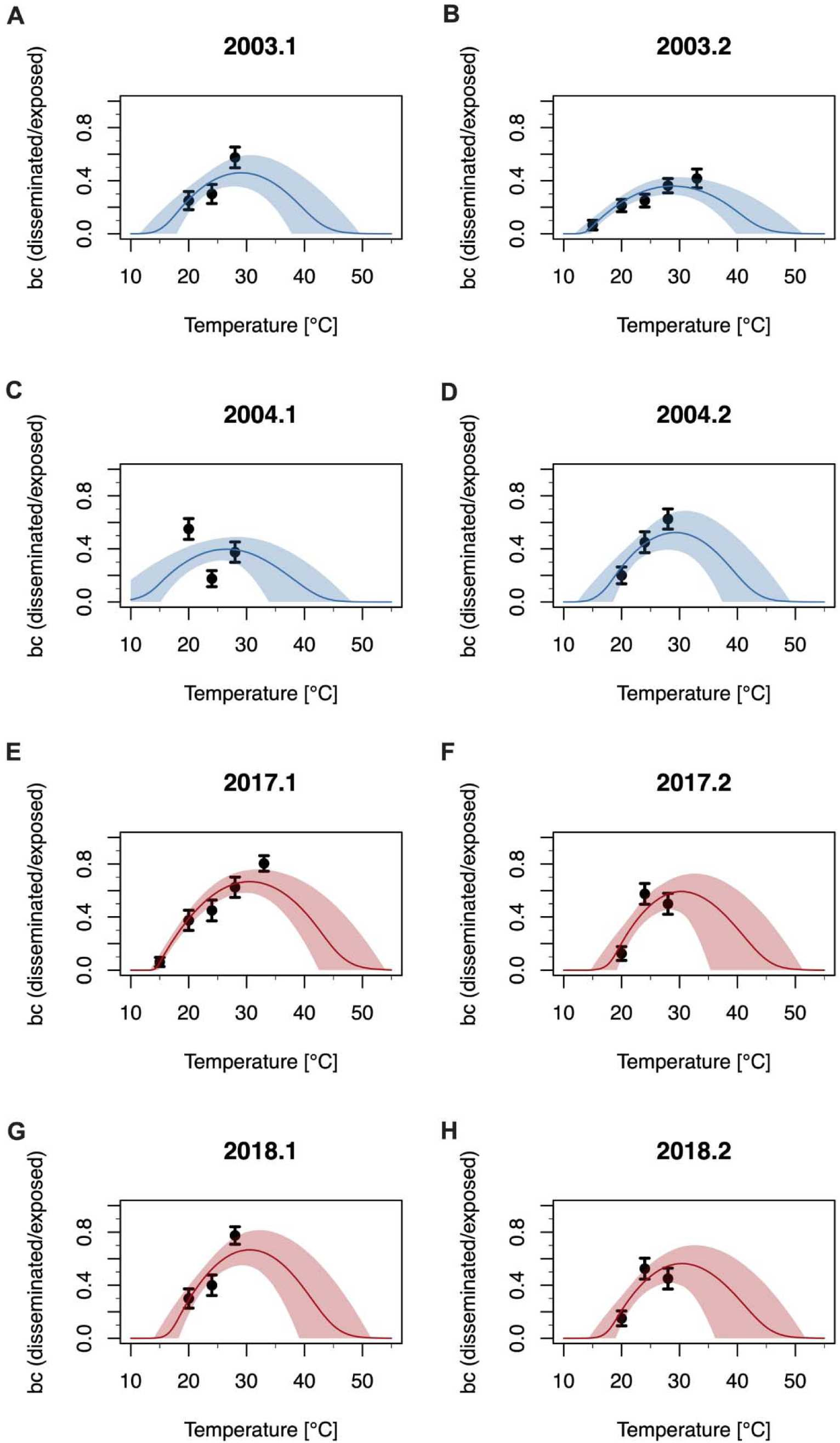
Thermal performance curves of *Culex pipiens* vector competence for historic and contemporary strains. (**A**) 2003.1, (**B**) 2003.2, (**C**) 2004.1, (**D**) 2004.2, (**E**) 2017.1, (**F**) 2017.2, (**G**) 2018.1, (**H**) 2018.2. Thermal response of vector competence *bc* (# disseminated/ # exposed) across temperature shown here. Solid lines are posterior means and shaded areas are 95% credible intervals. Historic strains are shown in blue and contemporary strains in red. Additional data points in B and E are from another experiment and are used to estimate the thermal limits of West Nile virus transmission.

**Figure S2.**
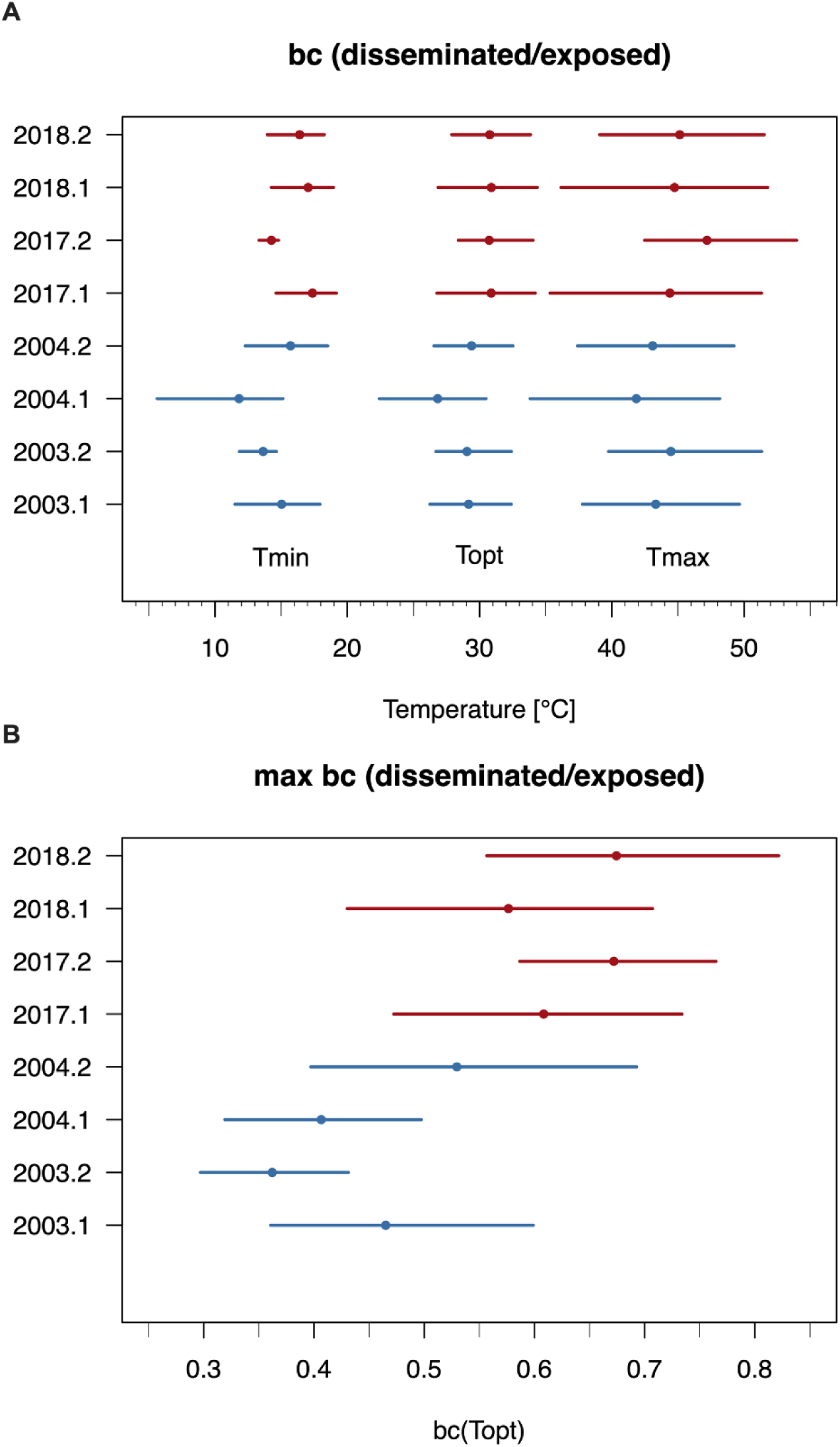
Estimated strain-specific TPC parameters for vector competence. **(A)** The estimated thermal minimum, optimum, and maximum of *Culex pipiens* WNV vector competence is shown for each strain. **(B)** The estimated *Culex pipiens* maximum WNV vector competence is shown for each strain. Points correspond to posterior means and lines to 95% credible intervals. Contemporary WNV strains are shown in red and historic strains are shown in blue.

**Figure S3.**
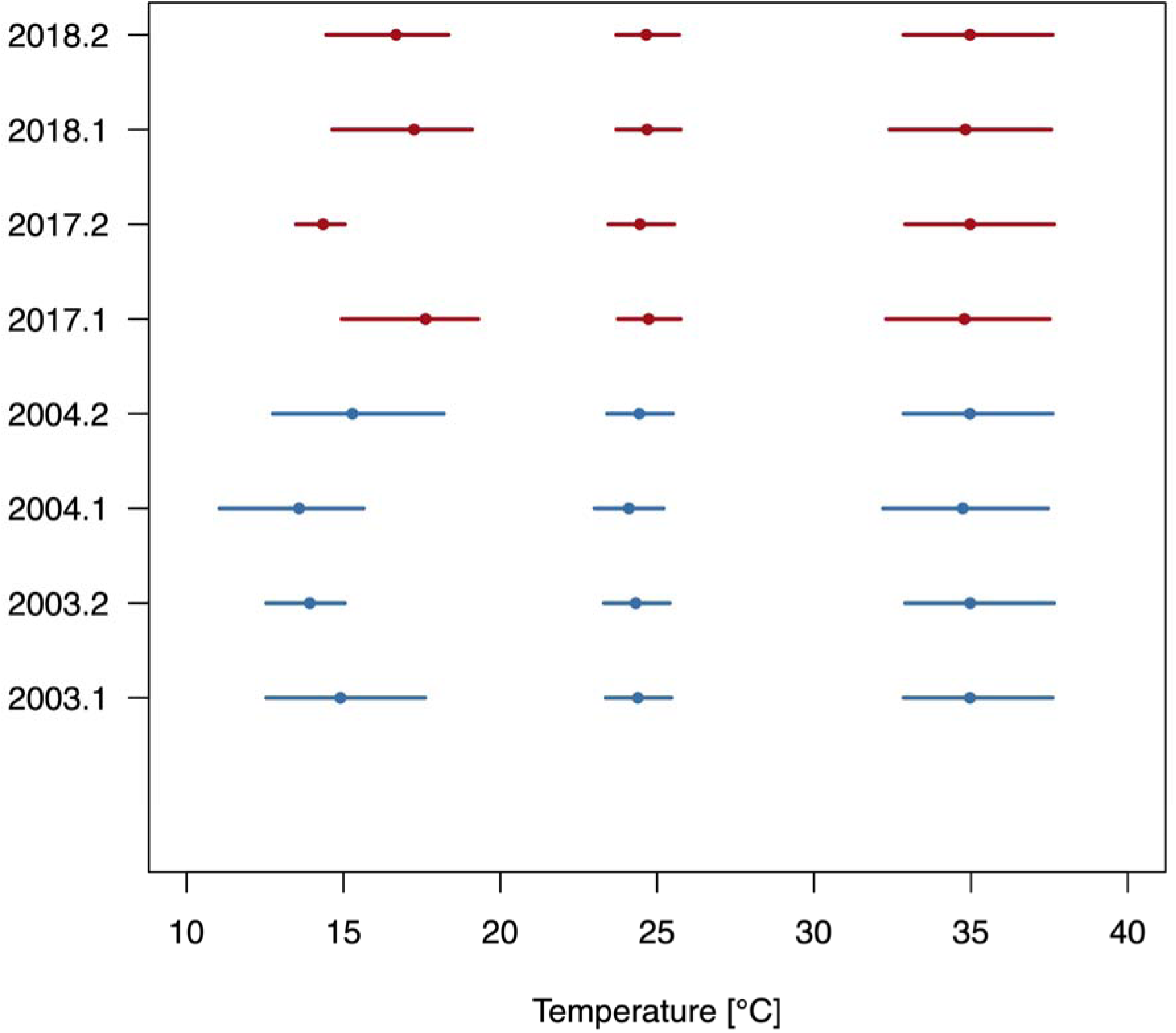
Estimated thermal optimum and limits for WNV transmission (relative R_0_) in *Cx pipiens*. Posterior mean and uncertainty estimates for the minimum, optimum, and maximum temperatures. Points show mean and lines show 95% credible intervals. Contemporary strains are shown in red and historic WNV strains are shown in blue.

**Figure S4.**
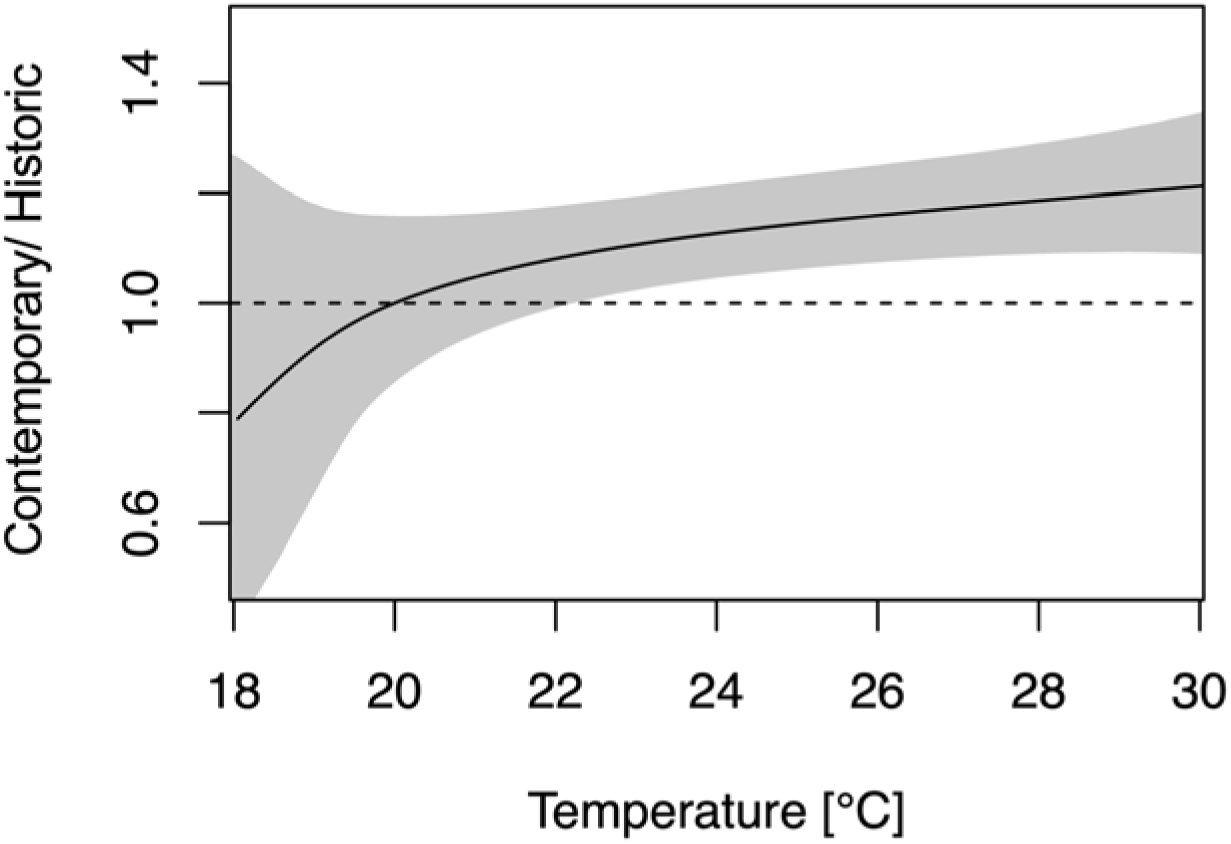
The ratio between the mean relative R_0_ of the four contemporary strains and the mean of the four historic strains is shown at each temperature for the TPC analysis. Contemporary strains have increased transmission on average than historic strains at high temperatures. Points correspond to posterior means and lines to 95% credible intervals. The line corresponds to the posterior mean and the shaded region to a 95% credible interval. A ratio of one (corresponding to equal transmission) is shown as a dotted line.

**Figure S5.**
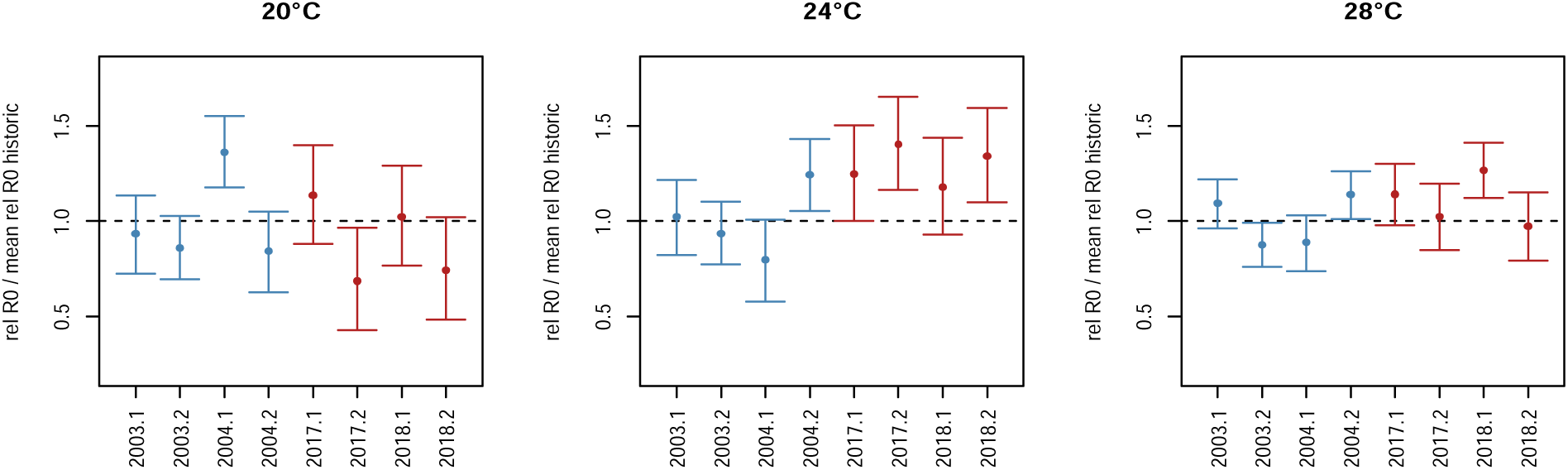
Transmission of individual WNV strains compared to the mean transmission of historic strains. The ratio between the relative R_0_ of a given strain and the mean relative R_0_ for historic strains from the point-wise analysis is shown at each temperature. Points correspond to posterior means and lines to 95% credible intervals. A ratio of one (corresponding to equal transmission to the historic mean) is shown as a dotted line.

## Acknowledgments

We thank the NYS Arbovirology Laboratory insectary staff for support and assistance. We thank the Wadsworth Center Media and Tissue Culture Facility for providing cells and media. ATC was supported by National Institutes of Health R01AI168097. EAM was supported by grants from the National Science Foundation (DEB-2011147 with Fogarty International Center), National Institutes of Health (R35GM133439, R01AI168097, R01AI102918), the Stanford Woods Institute for the Environment, King Center on Global Development, and Center for Innovation in Global Health.

