## Appendix for "Temperature influences West Nile virus evolution and adaptation"

#### Estimation of the temperature-dependence of $\mathcal{R}_0$

As in (Shocket et al 2020), we model the temperature-dependence of the basic reproductive number $\mathcal{R}_0$ for West Nile Virus as follows:

$$\text{relative } \mathcal{R}_0(T) = \sqrt{\frac{a^3(T)\mathrm{bc}(T)e^{-\frac{\mu(T)}{\mathrm{PDR}(T)}}\mathrm{EFGC}(T)\mathrm{EV}(T)p_{LA}(T)\mathrm{MDR}(T)}{\mu^3(T)}} \quad (1)$$

where $a(T)$ is the mosquito biting rate, $\mathrm{bc}(T)$ the vector competence of West Nile Virus, $\mu(T)$ the mosquito death rate (i.e. the inverse of lifespan), $\mathrm{PDR}(T)$ the pathogen development rate, $\mathrm{ER}(T)$ the eggs per raft laid by a female mosquito, $\mathrm{EFGC}(T)$ eggs per female per gonotrophic cycle, $p_{LA}(T)$ the proportion of mosquito larvae that survive to adulthood, $\mathrm{MDR}(T)$ the mosquito development rate (i.e. inverse of time for a larva to become an adult). Note that the expression for relative $\mathcal{R}_0(T)$ omits the number of hosts $N$ and the host recovery rate $r$ (which are assumed to be temperature independent) from the denominator. Thus, relative $\mathcal{R}_0(T)$ is proportional to the basic reproductive number $\mathcal{R}_0(T)$ and has the same temperature-dependence, but the proportionality constant is not known in the absence of these host parameters.

Estimates for vector competence $\mathrm{bc}(T)$ were obtained for the West Nile virus strains in this study, while posterior samples for the thermal performance curves of mosquitoes life history trait estimates were taken from Shocket et al 2020 (adapting the code published in this article to generate posterior samples). As described in the Methods, vector competence was estimated in two ways: a) a point-wise analysis, where vector competence and relative $\mathcal{R}_0$ were estimated only at the experimentally measured temperatures and b) a TPC analysis where both vector competence and relative $\mathcal{R}_0$ are estimated also for other temperatures. Models were fit with Bayesian methods using Markov Chain Monte Carlo (MCMC), as described in the Methods. To obtain posterior samples for relative $\mathcal{R}_0$, the parameter values corresponding to each iteration of the MCMC chains for each TPC in Shocket et al were used to calculate predicted trait values and were combined with posterior samples for vector competence. These predicted trait values were used to calculate posterior samples for relative $\mathcal{R}_0(T)$ with Equation 1.

We obtained posterior draws for relative $\mathcal{R}_0$ (Equation 1 in this Appendix) from the posterior draws of the thermal performance curves from individual traits. In the point-wise analysis, we do so only for the three temperatures that were measured for all strains ($20°\text{C}, 24°\text{C}, 28°\text{C}$). In the TPC analysis, we calculated the predicted trait values for each posterior draw in the MCMC chains for all traits in Equation 1 for a sequence of temperatures ranging from 10°C to 40°C, with a 0.025°C interval. The resulting trait values were used in Equation 1 to produce a posterior draw for relative $\mathcal{R}_0$. This procedure was repeated for each MCMC chain iteration to obtain samples from the posterior distribution of relative $\mathcal{R}_0$ at each temperature. The minimum, maximum and optimum temperatures for relative $\mathcal{R}_0$ were also calculated for each posterior sample in the TPC analyis: this allowed us to estimate the mean and 95% credible interval for these quantities.

### Point-wise analysis

In this section, we present results from the analysis shown in the main text, which is based on point estimates of vector competence at the specific temperatures that were measured experimentally. The statistical model used in the point-wise analysis makes minimal assumptions and makes conclusions limited to only the specific measured temperatures. The TPC analysis makes stronger modeling assumptions based on previous experiments, but provides estimates for transmission at other temperatures than those directly measured.

#### Statistical model

##### Likelihood

The total number of mosquitoes $n_{sT}$ that developed a disseminated infection from WNV strain $s$ at temperature $T$ was modeled as a binomial distribution

$$n_{sT}|\text{bc}_{sT} \sim \text{Binomial}(p = \text{bc}_{sT}, N = N_{sT})$$

where $N_{sT}$ is the total number of mosquitoes exposed to strain $s$ evaluated at temperature $T$ and $\text{bc}_{sT}$ is the probability that a mosquito of strain $s$ develops a disseminated West Nile virus infection at temperature $T$.

##### Prior distribution

A uniform prior distribution was used for vector competence for each strain and temperature.

$$\text{bc}_{sT} \sim \text{Uniform}(0, 1)$$

#### Posterior estimates for vector competence

The posterior estimates for vector competence $\text{bc}_{sT}$ (i.e. the proportion of exposed mosquitoes that develop a disseminated infection at temperature $T$) are shown below for each strain and temperature.

**Historic strains**

| Strain | Temperature | Mean | 2.5% | 97.5% | $\hat{R}$ | $n_{\text{eff}}$ |
|---|---|---|---|---|---|---|
| 2003.1 | 22 | 0.261 | 0.142 | 0.403 | 1.001 | 80000 |
| 2003.1 | 24 | 0.309 | 0.180 | 0.455 | 1.001 | 80000 |
| 2003.1 | 28 | 0.571 | 0.421 | 0.716 | 1.001 | 80000 |
| 2003.2 | 22 | 0.220 | 0.137 | 0.314 | 1.001 | 48000 |
| 2003.2 | 24 | 0.256 | 0.169 | 0.356 | 1.001 | 38000 |
| 2003.2 | 28 | 0.366 | 0.265 | 0.471 | 1.001 | 57000 |
| 2004.1 | 22 | 0.548 | 0.398 | 0.694 | 1.001 | 80000 |
| 2004.1 | 24 | 0.190 | 0.088 | 0.321 | 1.001 | 68000 |
| 2004.1 | 28 | 0.381 | 0.241 | 0.531 | 1.001 | 80000 |
| 2004.2 | 22 | 0.214 | 0.105 | 0.348 | 1.001 | 80000 |
| 2004.2 | 24 | 0.452 | 0.306 | 0.602 | 1.001 | 32000 |
| 2004.2 | 28 | 0.619 | 0.469 | 0.758 | 1.001 | 43000 |

**Contemporary strains**

| Strain | Temperature | Mean | 2.5% | 97.5% | $\hat{R}$ | $n_{\text{eff}}$ |
|---|---|---|---|---|---|---|
| 2017.1 | 22 | 0.381 | 0.242 | 0.531 | 1.001 | 80000 |
| 2017.1 | 24 | 0.453 | 0.307 | 0.603 | 1.001 | 31000 |
| 2017.1 | 28 | 0.619 | 0.470 | 0.759 | 1.001 | 32000 |
| 2017.2 | 22 | 0.143 | 0.056 | 0.262 | 1.001 | 54000 |
| 2017.2 | 24 | 0.571 | 0.421 | 0.715 | 1.001 | 52000 |
| 2017.2 | 28 | 0.500 | 0.352 | 0.648 | 1.001 | 80000 |
| 2018.1 | 22 | 0.310 | 0.181 | 0.456 | 1.001 | 49000 |
| 2018.1 | 24 | 0.405 | 0.264 | 0.555 | 1.001 | 80000 |
| 2018.1 | 28 | 0.762 | 0.624 | 0.877 | 1.001 | 80000 |
| 2018.2 | 22 | 0.166 | 0.072 | 0.291 | 1.001 | 80000 |
| 2018.2 | 24 | 0.523 | 0.373 | 0.671 | 1.001 | 71000 |
| 2018.2 | 28 | 0.452 | 0.306 | 0.602 | 1.001 | 72000 |

**Relative $\mathcal{R}_0$**

The posterior estimates for relative $\mathcal{R}_0$ for each strain are shown below:

| group | strain | 20°C | | | 24°C | | | 28°C | | |
|---|---|---|---|---|---|---|---|---|---|---|
| | | Mean | 2.5% | 97.5% | Mean | 2.5% | 97.5% | Mean | 2.5% | 97.5% |
| historic | 2003.1 | 46.16 | 26.40 | 71.94 | 67.04 | 43.13 | 96.45 | 61.76 | 37.70 | 92.07 |
| historic | 2003.2 | 42.40 | 25.02 | 64.55 | 61.18 | 40.62 | 86.64 | 49.39 | 29.94 | 74.01 |
| historic | 2004.1 | 67.14 | 40.88 | 100.53 | 52.27 | 31.25 | 78.72 | 50.28 | 29.87 | 76.58 |
| historic | 2004.2 | 41.63 | 23.18 | 65.78 | 81.35 | 54.31 | 114.13 | 64.32 | 39.37 | 95.69 |
| contemporary | 2017.1 | 55.83 | 33.17 | 84.72 | 81.39 | 54.41 | 114.29 | 64.32 | 39.39 | 95.64 |
| contemporary | 2017.2 | 33.73 | 17.48 | 55.65 | 91.57 | 62.32 | 126.80 | 57.72 | 34.86 | 86.40 |
| contemporary | 2018.1 | 50.29 | 29.33 | 77.27 | 76.92 | 51.03 | 108.84 | 71.43 | 44.15 | 105.28 |
| contemporary | 2018.2 | 36.56 | 19.51 | 59.36 | 87.61 | 59.29 | 121.90 | 54.86 | 32.99 | 82.69 |

Note that these relative $\mathcal{R}_0$ values are not interpretable in an absolute sense, as scaling them to a full $\mathcal{R}_0$ would require knowing the number of hosts and the host recovery rate, which are unknown (see Shocket et al for details). For example, a relative $\mathcal{R}_0$ of 1 does not hold any particular meaning (unlike for a full $\mathcal{R}_0$, for which it indicates transmission is high enough for an epidemic to occur). However, they are comparable in a relative sense. That is, larger values correspond to more transmission both across temperatures within a strain, and when comparing different strains (with the assumption that the number of hosts and recovery rate are equal among the strains).

##### $\mathcal{R}_0$ ratio

The mean relative $\mathcal{R}_0$ for strains in group $g$ at temperature $T$ is given by

$$M_{GT} = \frac{1}{n_G} \sum_{s=1}^{n_g} \text{relative } \mathcal{R}_{0s,T}$$

where $g$ corresponds to a group of strains (i.e. contemporary or historic) and $n_G$ to the number of strains in that group. We used the ratio $\rho_T$ of the mean relative $\mathcal{R}_0$ of the contemporary and historic strains

$$\rho_T = \frac{M_{CT}}{M_{HT}}$$

as a measure of the relative transmission of contemporary and historic strains at temperature $T$, where a ratio of one corresponds to equal transmission between historic and contemporary strains.

The posterior estimates of $\rho_T$ at each measured temperature are as follows:

| Temperature | Mean | 2.5% | 97.5% | $\Pr(\rho_T > 1)$ |
|---|---|---|---|---|
| 20 | 0.897 | 0.747 | 1.062 | 0.1023 |
| 24 | 1.293 | 1.128 | 1.480 | 0.9999 |
| 28 | 1.102 | 0.997 | 1.215 | 0.9710 |

Our results show that transmission for the contemporary WNV strains studied in this work is higher on average than for the historic WNV strains at 24°C and 28°C, with the largest difference ocurring at 24°C.

##### Individual strains $\mathcal{R}_0$ ratio

We also compared the transmission of each individual strain relative to the mean of the four historic strains through the ratio

$$\rho_{sT} = \frac{\text{relative } \mathcal{R}_{0,s}}{M_{HT}}$$

where a ratio of one corresponds to equal transmission relative to the historic mean at that temperature, which is more interpretable than the relative $\mathcal{R}_0$ values. As various sources of uncertainty for relative $\mathcal{R}_0$ are common to all strains (e.g. mosquito life history traits independent of WNV infection), this ratio also has reduced uncertainty compared to the relative $\mathcal{R}_0$ values. The posterior distribution of this ratio is summarized below.

| group | strain | 20°C | | | 24°C | | | 28°C | | |
|---|---|---|---|---|---|---|---|---|---|---|
| | | Mean | 2.5% | 97.5% | Mean | 2.5% | 97.5% | Mean | 2.5% | 97.5% |
| historic | 2003.1 | 0.935 | 0.727 | 1.136 | 1.024 | 0.823 | 1.217 | 1.095 | 0.962 | 1.220 |
| historic | 2003.2 | 0.860 | 0.698 | 1.027 | 0.936 | 0.773 | 1.103 | 0.875 | 0.760 | 0.990 |
| historic | 2004.1 | 1.363 | 1.177 | 1.551 | 0.797 | 0.580 | 1.008 | 0.890 | 0.739 | 1.030 |
| historic | 2004.2 | 0.843 | 0.629 | 1.050 | 1.244 | 1.055 | 1.432 | 1.140 | 1.013 | 1.262 |
| contemporary | 2017.1 | 1.135 | 0.884 | 1.398 | 1.247 | 1.000 | 1.504 | 1.141 | 0.978 | 1.301 |
| contemporary | 2017.2 | 0.686 | 0.429 | 0.969 | 1.403 | 1.165 | 1.652 | 1.024 | 0.849 | 1.196 |
| contemporary | 2018.1 | 1.022 | 0.768 | 1.290 | 1.179 | 0.929 | 1.441 | 1.268 | 1.121 | 1.412 |
| contemporary | 2018.2 | 0.743 | 0.485 | 1.022 | 1.343 | 1.101 | 1.594 | 0.974 | 0.792 | 1.150 |

### Thermal performance curve analysis

As a complementary analysis to the results shown in the main text, which are based on point estimates of vector competence at specific temperatures measured experimentally, we estimated thermal performance curves (TPCs) for vector competence for the individual historic and contemporary West Nile Virus strains studied here and used these curves to estimate the effects on this variation in transmission. To do this, we followed the approach in [1], where TPCs for various mosquito and virus traits are combined in a mechanistic temperature-dependent $\mathcal{R}_0$ model of West Nile Virus transmission. This allowed us to explore the possible consequences of strain-specific differences in vector competence in transmission at other temperatures than those measured directly.

One limitation of this analysis is that estimating TPCs is challenging when there is a limited measured temperature range, as is the case for many of the strains here. We took the following steps to mitigate this issue:

1. We include vector competence data at two additional temperatures from a separate experiment for two strains (see below).

2. Rather than fitting the TPCs for each strain individually, we took a hierarchical modeling approach. This made it possible to improve the estimated TPCs by performing partial pooling of data from different strains.

3. We used informative prior distributions based on previous fits to vector competence data of other WNV strains in *Culex pipiens* (the same mosquito species of interest in this work) on the TPC parameters. This was especially important to constrain the thermal limits to reasonable ranges in cases where there is no measured data near temperature extremes.

While these steps were helpful in improving the TPC estimates, the estimated curves and the underlying TPC parameters (such as the optimal temperature and thermal limits) still had high uncertainty in many cases. Because of this, our analysis is only able to conclusively detect relatively large differences in the TPC parameters among the strains and smaller differences may remain undetected. In some cases, our analysis suggested differences may exist among the strains, but the statistical evidence was not strong enough for the result to be conclusive (which we operationally define here as a posterior probability greater than or equal to 0.95 that the result is true). While these results need independent confirmation, we report them here as they can be helpful for generating hypotheses for future studies that measure a wider range of temperatures.

It must also be noted that the reported credible intervals and posterior probabilities depend on the accuracy of the modeling assumptions, which include that the assumed quadriatic TPC functional form is a good approximation of the true shape of the vector competence TPC for the WNV strains studied here. While it is difficult to check this assumption directly from the data in cases where there are only a few measured temperatures, we believe the quadratic model is reasonable since it has been previously found to be a good fit to vector competence data in studies with other WNV strains [1], and it seems to provide a good fit to strains in this work for which more temperature measurements are available. Future studies with measurements at more temperatures will make it possible to evaluate this assumption directly with the experimental data.

#### Additional data for vector competence TPC analysis

In a separate experiment to that described in the main text, data for infection and dissemination at two additional temperatures was obtained for one historic and one contemporary West Nile Virus strain, shown below.

| Strain | Temperature | Infection $n$ | (%) | Dissemination $n$ | (%) | Total |
|---|---|---|---|---|---|---|
| 2003.2 | 15°C | 6 | 13.04% | 3 | 6.52% | 46 |
| 2003.2 | 33°C | 35 | 72.92% | 20 | 41.67% | 48 |
| 2017.1 | 15°C | 13 | 26.53% | 3 | 6.12% | 49 |
| 2017.1 | 33°C | 42 | 91.30% | 37 | 80.43% | 46 |

This additional data was used in the thermal performance curve fits for the corresponding strains (i.e. for these strains there were mesaurements at five different temperatures available, rather than three as for the rest of the strains).

#### Thermal performance curves for vector competence

As in previous work [1], the dissemination proportion was modeled with a quadratic model, given by

$$\mathrm{bc}(T) = c(T - T_{\mathrm{min}})(T_{\mathrm{max}} - T)$$

where $T_{\mathrm{min}}$ and $T_{\mathrm{max}}$ are the minimum and maximum temperatures and $c > 0$ is an arbitrary constant. In the quadratic model, the optimal temperature is given by

$$T_{\text{opt}} = \frac{T_{\min} + T_{\max}}{2}$$

This result can be used to reparametrize this model in terms of the maximum dissemination proportion $\text{bc}_{\max}$ rather than the constant $c$, since

$$\text{bc}_{\max} = c(T_{\text{opt}} - T_{\min})(T_{\max} - T_{\text{opt}})$$

which, after substitution and some algebra yields

$$c = \frac{4\text{bc}_{\max}}{(T_{\max} - T_{\min})^2}$$

This expression can be substituted back into the equation for $\text{bc}(T)$ to yield the quadratic model reparametrized in terms of the maximum dissemination proportion

$$\text{bc}(T) = \frac{4\text{bc}_{\max}}{(T_{\max} - T_{\min})^2} (T - T_{\min})(T_{\max} - T)$$

**Statistical model**

**Likelihood**

The total number of disseminated mosquitoes $n_{is}$ from WNV strain $s$ at temperature $T_{is}$ was modeled as a binomial distribution where the probability $\text{bc}(T)$ of a mosquito developing a disseminated West Nile virus infection follows a quadratic thermal performance curve

$$n_{is}|T_{is}, N_{is}, \mathcal{P}_s \sim \text{Binomial}(p = \text{bc}(T_{is}; \mathcal{P}_s), N = N_{is})$$

where $N_{is}$ is the total number of mosquitoes evaluated at temperature $T_{is}$ and $\mathcal{P}_s = \{T_{\min,s}, T_{\max,s}, \text{bc}_{\max,s}\}$ are the parameters for the quadratic TPC model corresponding to strain $s$. A hierarchical model was employed for each TPC parameter, with separate hierarchical models used for historical (2003-2004) and contemporary (2017-2018) strains, defined as follows:

$$\begin{aligned} T_{\min,s}|T_{\min,g}, \sigma_{T_{\min,g}} &\sim \text{Normal}(\mu = T_{\min,g}, \sigma = \sigma_{T_{\min,g}}) \\ T_{\max,s}|T_{\max,g}, \sigma_{T_{\max,g}} &\sim \text{Normal}(\mu = T_{\max,g}, \sigma = \sigma_{T_{\max,g}}) \\ \text{bc}_{\max,s}|\text{bc}_{\max,g}, v_{r_{\max,g}} &\sim \text{Beta}(\mu = \text{bc}_{\max,g}, v = v_{r_{\max,g}}) \end{aligned}$$

where $\mathcal{P}_g = \{T_{\min,g}, T_{\max,g}, \text{bc}_{\max,g}\}$ are the grand means for each TPC parameter for strains of group $g$ (where $g \in \{\text{historic, contemporary}\}$). Note that the Beta distribution is parametrized in terms of the mean $\mu = \frac{\alpha}{\alpha+\beta}$ and sample size $v = \alpha + \beta$.

##### Prior distributions

Informative prior distributions were used for the grand means of the thermal performance curve parameters $T_{\min,g}$, $T_{\max,g}$ and $\text{bc}_{\max,g}$, based on previous fits to dissemination proportions in [1]. The specific priors used were

$$\begin{aligned} T_{\min,g} &\sim \text{Normal}(\mu = 16.8, \sigma = 2) \\ T_{\max,g} &\sim \text{Normal}(\mu = 38.9, \sigma = 3) \\ \text{bc}_{\max,g} &\sim \text{Beta}(\mu = 0.35, v = 4) \end{aligned}$$

The means for $T_{\min,g}$ and $T_{\max,g}$ were chosen to match the posterior mean estimates from Shocket et al, and the priors were given inflated variances to account for the possibility of the strains studied here differing from those studied previously. As this previous study did not parametrize the quadratic model in terms of the maximum dissemination proportion, no estimates are provided for this quantity explicitly. However, the posterior mean and 95% credible interval for $\text{bc}_{\max}$ was approximated visually from the plot in Shocket et al.

The posterior means and credible intervals from Shocket et al and the 95% prior CI for the grand mean of the TPC parameters in this work are summarized in the table below (where NE is used to indicate that a parameter was not estimated in Shocket et al and was estimated visually):

| Parameter | Shocket mean | Shocket 95% CI | Prior 95% CI |
|---|---|---|---|
| $T_{\min}$ | 16.8 | $(15.0, 17.9)$ | $(12.9, 20.7)$ |
| $T_{\max}$ | 38.9 | $(36.1, 44.1)$ | $(33.0, 44.8)$ |
| $\text{bc}_{\max}$ | $\approx 0.35$ (NE) | $\approx (0.25, 0.45)$ (NE) | $(0.03, 0.80)$ |

The following weakly informative prior distributions were used for the dispersion parameters of the hierarchical model:

$$\begin{aligned} \sigma_{T_{\min,g}} &\sim \text{HalfNormal}(\mu = 0, \sigma = 2) \\ \sigma_{T_{\max,g}} &\sim \text{HalfNormal}(\mu = 0, \sigma = 2) \\ \frac{1}{\sqrt{v_{r_{\max},g}}} &\sim \text{HalfNormal}(\mu = 0, \sigma = 1) \end{aligned}$$

Small values of these hyperparameters correspond to more "partial pooling" of the observations of the different strains to estimate the corresponding TPC parameter for each strain, while large values correspond to little to no pooling (and are equivalent to fitting a separate model for each strain).

A HalfNormal distribution is a Normal distribution truncated to only have nonzero probability density for values at or above the peak of the distribution (i.e. in the cases above, where $\mu = 0$, a halfNormal distribution only has support for nonnegative values). The priors for $\sigma_{T_{\min,g}}$ and $\sigma_{T_{\max,g}}$ correspond to the assumption that the standard deviation for the minimum and maximum temperatures for WNV dissemination of the various historical (resp. contemporary) strains is approxmately 95% likely *a priori* to be in the interval $[0°\text{C}, 4°\text{C}]$. In other words, this prior assumes that it is unlikely for a "typical" difference between a strain's $T_{\text{min}}$ (resp. $T_{\text{max}}$) and the mean $T_{\text{min}}$ (resp. $T_{\text{max}}$) across all strains of the corresponding group (historic or contemporary) is more than 4°C. The transformation $\frac{1}{\sqrt{v_{r_{\text{max}},g}}}$ was used for the sample size hyperparameter as the population standard deviation of the Beta distribution scales with respect to the sample size as $\frac{1}{\sqrt{v_{r_{\text{max}},g}}}$. The prior for $\frac{1}{\sqrt{v_{r_{\text{max}},g}}}$ corresponds to a weakly informative prior for a parameter in the unit scale.

#### Posterior estimates

The posterior estimates for all model parameters are shown below for each strain. The hyperparameters for the historic and contemporary strains (i.e. grand mean and standard deviation/sample size) for each TPC parameter are indicated with an H and C, respectively. The optimal temperature for each strain was estimated from the posterior draws, through the equation $T_{\text{opt}} = \frac{T_{\text{min}}+T_{\text{max}}}{2}$ which relates the optimum temperature to the minimum and maximum temperatures in the quadratic model.

##### Historic strains

##### Minimum temperature

| Strain | Parameter | Mean | 2.5% | 97.5% | $\hat{R}$ | $n_{\text{eff}}$ |
|---|---|---|---|---|---|---|
| H | $T_{\text{min},g}$ | 14.7 | 12.4 | 17.3 | 1.0011 | 22000 |
| H | $\sigma_{T_{\text{min},g}}$ | 2.1 | 0.1 | 4.8 | 1.0010 | 80000 |
| 2003.1 | $T_{\text{min},s}$ | 15.0 | 11.5 | 17.9 | 1.0010 | 80000 |
| 2003.2 | $T_{\text{min},s}$ | 13.6 | 11.8 | 14.6 | 1.0010 | 64000 |
| 2004.1 | $T_{\text{min},s}$ | 11.8 | 5.6 | 15.1 | 1.0010 | 53000 |
| 2004.2 | $T_{\text{min},s}$ | 15.7 | 12.3 | 18.5 | 1.0010 | 62000 |

##### Maximum temperature

| Strain | Parameter | Mean | 2.5% | 97.5% | $\hat{R}$ | $n_{\text{eff}}$ |
|---|---|---|---|---|---|---|
| H | $T_{\text{max},g}$ | 42.7 | 38.2 | 47.3 | 1.0010 | 36000 |
| H | $\sigma_{T_{\text{max},g}}$ | 2.0 | 0.1 | 5.2 | 1.0011 | 15000 |
| 2003.1 | $T_{\text{max},s}$ | 43.3 | 37.8 | 49.6 | 1.0011 | 24000 |
| 2003.2 | $T_{\text{max},s}$ | 44.5 | 39.7 | 51.3 | 1.0010 | 80000 |
| 2004.1 | $T_{\text{max},s}$ | 41.8 | 33.8 | 48.2 | 1.0010 | 37000 |
| 2004.2 | $T_{\text{max},s}$ | 43.1 | 37.4 | 49.2 | 1.0010 | 80000 |

**Maximum dissemination proportion**

| Strain | Parameter | Mean | 2.5% | 97.5% | $\hat{R}$ | $n_{\text{eff}}$ |
|---|---|---|---|---|---|---|
| H | $\text{bc}_{\max,g}$ | 0.437 | 0.307 | 0.583 | 1.0011 | 32000 |
| H | $\frac{1}{\sqrt{v_{r_{\max,g}}}}$ | 0.249 | 0.032 | 0.672 | 1.0011 | 20000 |
| 2003.1 | $\text{bc}_{\max,s}$ | 0.465 | 0.361 | 0.599 | 1.0011 | 29000 |
| 2003.2 | $\text{bc}_{\max,s}$ | 0.362 | 0.297 | 0.431 | 1.0010 | 80000 |
| 2004.1 | $\text{bc}_{\max,s}$ | 0.407 | 0.319 | 0.497 | 1.0010 | 42000 |
| 2004.2 | $\text{bc}_{\max,s}$ | 0.530 | 0.397 | 0.693 | 1.0011 | 28000 |

**Optimum temperature**

| Strain | Parameter | Mean | 2.5% | 97.5% | $\hat{R}$ | $n_{\text{eff}}$ |
|---|---|---|---|---|---|---|
| 2003.1 | $T_{\text{opt},s}$ | 29.2 | 26.2 | 32.4 | 1.0011 | 26000 |
| 2003.2 | $T_{\text{opt},s}$ | 29.0 | 26.7 | 32.4 | 1.0010 | 80000 |
| 2004.1 | $T_{\text{opt},s}$ | 26.8 | 22.4 | 30.5 | 1.0011 | 29000 |
| 2004.2 | $T_{\text{opt},s}$ | 29.4 | 26.5 | 32.5 | 1.0010 | 56000 |

**Contemporary strains**

**Minimum temperature**

| Strain | Parameter | Mean | 2.5% | 97.5% | $\hat{R}$ | $n_{\text{eff}}$ |
|---|---|---|---|---|---|---|
| C | $T_{\min,g}$ | 16.3 | 14.4 | 18.4 | 1.0010 | 37000 |
| C | $\sigma_{T_{\min,g}}$ | 1.9 | 0.6 | 3.7 | 1.0011 | 21000 |
| 2017.2 | $T_{\min,s}$ | 17.4 | 14.6 | 19.2 | 1.0013 | 7500 |
| 2017.1 | $T_{\min,s}$ | 14.3 | 13.3 | 14.8 | 1.0010 | 80000 |
| 2018.2 | $T_{\min,s}$ | 17.0 | 14.3 | 19.0 | 1.0011 | 19000 |
| 2018.1 | $T_{\min,s}$ | 16.4 | 14.0 | 18.3 | 1.0010 | 36000 |

**Maximum temperature**

| Strain | Parameter | Mean | 2.5% | 97.5% | $\hat{R}$ | $n_{\text{eff}}$ |
|---|---|---|---|---|---|---|
| C | $T_{\max,g}$ | 44.4 | 39.3 | 49.3 | 1.0010 | 55000 |
| C | $\sigma_{T_{\max,g}}$ | 2.3 | 0.1 | 5.6 | 1.0011 | 15000 |
| 2017.2 | $T_{\max,s}$ | 44.4 | 35.3 | 51.3 | 1.0010 | 80000 |
| 2017.1 | $T_{\max,s}$ | 47.2 | 42.5 | 54.0 | 1.0014 | 6600 |
| 2018.2 | $T_{\max,s}$ | 44.7 | 36.2 | 51.8 | 1.0010 | 80000 |
| 2018.1 | $T_{\max,s}$ | 45.1 | 39.1 | 51.5 | 1.0011 | 18000 |

**Maximum dissemination proportion**

| Strain | Parameter | Mean | 2.5% | 97.5% | $\hat{R}$ | $n_{\text{eff}}$ |
|---|---|---|---|---|---|---|
| C | $\text{bc}_{\max,g}$ | 0.614 | 0.443 | 0.732 | 1.0010 | 80000 |
| C | $\frac{1}{\sqrt{v_{r_{\max},g}}}$ | 0.214 | 0.010 | 0.667 | 1.0011 | 16000 |
| 2017.2 | $\text{bc}_{\max,s}$ | 0.608 | 0.472 | 0.734 | 1.0011 | 25000 |
| 2017.1 | $\text{bc}_{\max,s}$ | 0.672 | 0.587 | 0.765 | 1.0010 | 65000 |
| 2018.2 | $\text{bc}_{\max,s}$ | 0.576 | 0.430 | 0.707 | 1.0011 | 24000 |
| 2018.1 | $\text{bc}_{\max,s}$ | 0.674 | 0.557 | 0.822 | 1.0010 | 70000 |

**Optimum temperature**

| Strain | Parameter | Mean | 2.5% | 97.5% | $\hat{R}$ | $n_{\text{eff}}$ |
|---|---|---|---|---|---|---|
| 2017.2 | $T_{\text{opt},s}$ | 30.9 | 26.8 | 34.2 | 1.0010 | 80000 |
| 2017.1 | $T_{\text{opt},s}$ | 30.7 | 28.4 | 34.0 | 1.0014 | 6600 |
| 2018.2 | $T_{\text{opt},s}$ | 30.9 | 26.9 | 34.4 | 1.0010 | 80000 |
| 2018.1 | $T_{\text{opt},s}$ | 30.8 | 27.9 | 33.9 | 1.0011 | 25000 |

**Changes in mean peak dissemination and thermal optima**

As a direct way to test whether contemporary strains have increased maximum dissemination proportions on average compared to historic strains, we obtain the posterior distribution for the statistic

$$\Delta_{\text{bc}_{\max}} = \text{bc}_{\max,\text{contemporary}} - \text{bc}_{\max,\text{historic}}$$

where $\text{bc}_{\max,g}$ is the grand mean for the maximum dissemination proportion in the hierarchical model for the contemporary and historic group, respectively, and calculate the posterior probability that $\Delta_{\text{bc}_{\max}} > 0$.

As the thermal optimum is not directly a model parameter in the quadratic model, but is a derived quantity calculated from the minimum and maximum temperatures, we define the mean optimal temperature for group $g$ (either historic or contemporary) as

$$T_{\text{opt},g} = \frac{1}{n_g} \sum_{s=1}^{n_g} T_{\text{opt},s}$$

where $T_{\text{opt},s}$ is the optimal temperature for strain $s$ and $n_g$ is the number of strains in group $g$ (either contemporary or historic). Using this definition, we obtain the posterior distribution for the mean optimal temperature for the historic and contemporary strains from the posterior samples for the $T_{\text{opt},s}$ and calculate the statistic

$$\Delta_{T_{\text{opt}}} = T_{\text{opt,contemporary}} - T_{\text{opt,historic}}$$

to test whether contemporary strains have an increased optimum temperature compared to historic strains by evaluating the posterior probability that $\Delta_{T_{\text{opt}}} > 0$.

| $\theta$ | Mean | 2.5% | 97.5% | $\Pr(\theta > 0)$ |
|---|---|---|---|---|
| $\Delta_{\text{bc}_{\text{max}}}$ | 0.177 | -0.038 | 0.354 | 0.955 |
| $\Delta_{T_{\text{opt}}}$ | 2.2 | -1.0 | 5.4 | 0.912 |

Our results show that, on average, contemporary strains have increased peak vector competence compared to historic strains. Our analysis also suggests that contemporary strains may have a higher thermal optimum for dissemination compared to historic strains, although this result is not conclusive due to the high uncertainty on the estimated thermal optima. Additional studies where vector competence for individual strains is measured at a wider range of temperatures would make it possible to better quantify this possible difference in thermal optima between historic and contemporary strains.
